# Selective inhibition of hsp90 paralogs: Uncovering the role of helix 1 in Grp94-selective ligand binding

**DOI:** 10.1101/2023.07.31.551342

**Authors:** Nanette L. S. Que, Paul M. Seidler, Wen J. Aw, Gabriela Chiosis, Daniel T. Gewirth

**Author notes:** To whom correspondence should be addressed: Daniel T. Gewirth, Hauptman Woodward Medical Research Institute 700 Ellicott Street, Buffalo, NY 14203 USA.

## Abstract

Grp94 is the endoplasmic reticulum paralog of the hsp90 family of chaperones, which have been targeted for therapeutic intervention via their highly conserved ATP binding sites. The design of paralog-selective inhibitors relies on understanding the protein structural elements that drive higher affinity in selective inhibitors. Here, we determined the structures of Grp94 and Hsp90 in complex with the Grp94-selective inhibitor PU-H36, and of Grp94 with the non-selective inhibitor PU-H71. In Grp94, PU-H36 derives its higher affinity by utilizing Site 2, a Grp94-specific side pocket adjoining the ATP binding cavity, but in Hsp90 PU-H36 occupies Site 1, a side pocket that is accessible in all paralogs with which it makes lower affinity interactions. The structure of Grp94 in complex with PU-H71 shows only Site 1 binding. While changes in the conformation of helices 4 and 5 in the N-terminal domain occur when ligands bind to Site 1 of both Hsp90 and Grp94, large conformational shifts that also involve helix 1 are associated with the engagement of the Site 2 pocket in Grp94 only. Site 2 in Hsp90 is blocked and its helix 1 conformation is insensitive to ligand binding. To understand the role of helix 1 in ligand selectivity, we tested the binding of PU-H36 and other Grp94-selective ligands to chimeric Grp94/Hsp90 constructs. These studies show that helix 1 is the major determinant of selectivity for Site 2 targeted ligands, and also influences the rate of ATPase activity in Hsp90 paralogs.

## Introduction

The hsp90 chaperones are responsible for the regulation and maturation of a diverse set of proteins that contribute to the maintenance of cell homeostasis ^1^. The client proteins of these chaperones include soluble and membrane-bound proteins, transcription factors, regulatory and checkpoint kinases, growth factors, angiogenesis promoters, metalloproteinases, and telomerases ^2–4^. Inhibition or knock-downs of hsp90 chaperones leads to the loss of function of their clients. Because the clients of hsp90s include key players in cancer, neurodegenerative and inflammatory diseases, as well as in innate immunity, the chaperones have been targeted for therapeutic intervention ^5–9^.

Higher eukaryotes contain four hsp90 paralogs in three compartments: Hsp90α and Hsp90β (cytosol), Grp94 (endoplasmic reticulum, ER), and Trap-1 (mitochondria). All of the paralogs share a common organization, with an N-terminal ATP binding and regulatory domain (NTD), a middle domain, and a C-terminal dimerization domain. Chaperone function is driven by conformational changes that are associated with ATP binding to the NTD and its subsequent hydrolysis ^3,5,10–12^. Small molecule inhibitors that disrupt nucleotide binding prevent the functionally relevant conformational changes and result in abortive client maturation. To date, hsp90 inhibitors have incorporated at least 19 different chemical scaffolds with a number of these inhibitors progressing into clinical trials ^13,14^. The first generation of these molecules were pan-hsp90 inhibitors that targeted all four paralogs, but with the failure to secure FDA approval for these drug candidates, subsequent efforts have evolved to include the targeting of individual paralogs ^9,15–21^.

The NTDs of the four hsp90 paralogs exhibit sequence identities of 50% or more, and within the ATP binding cavity the identity exceeds 70%. Despite the challenge posed by this high sequence conservation for the design of paralog-selective inhibitors, early efforts that focused on screening of chemical libraries identified inhibitors with paralog selective binding properties. Co-crystal structures with these inhibitors revealed that their binding is multipartite, where insertion of a ligand in the central ATP cavity is paired with occupancy of three adjoining pockets, termed Site 1, Site 2, and Site 3 (**Figure 1A**) ^5^. The ability of a ligand to access and stably bind to these compound sites in one paralog but not another contributes to binding selectivity.

**FIGURE 1.**
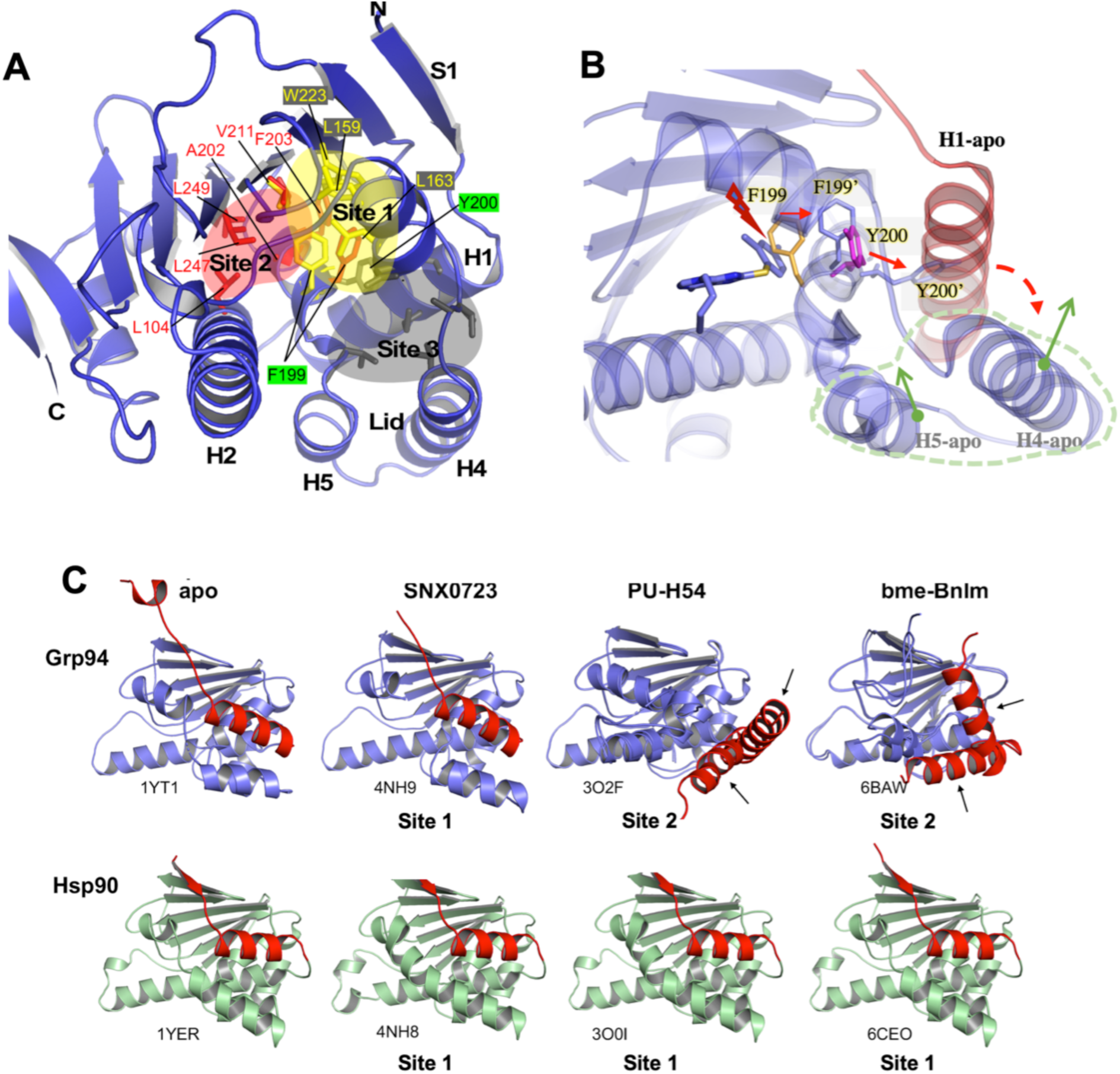
Inhibitors binding in the Grp94 ATP binding cavity access Sites 1, 2, and 3. (A) Residues comprising Sites 1 (yellow), 2 (red), and 3 (gray) mapped onto the Grp94 structure. (B) Cascade of conformational changes in the lid of Grp94 when Site 2 binding occurs. Binding of PU-H54 (blue) with the 8-aryl moiety in Site 2 of apo Grp94 (PDB 1YT1) forces the side chain of Phe199 (orange) away from the pocket to a new position F199’ (blue) to relieve a steric clash (red lightning symbol) with the 8-aryl moiety of the ligand. Movement of the Phe199 side chain impacts the position of Tyr200 (magenta) and results in conformational changes to the lid subdomain, particularly S1/H1. (C) Examples of Grp94 and Hsp90 S1/H1 conformations as a function of bound ligand. The 2 monomers in the asymmetric unit of 3O2F and 6BAW are shown superimposed; arrows denote alternate conformations. Compared to binding in Sites 1 and 3, binding in Site 2 induces large conformational changes in the lid of Grp94 but not Hsp90.

Sites 1, 2 and 3 are occluded in the apo form of the protein, and access depends on the ability of the NTD to undergo conformational changes in the “lid” subdomain, comprised of helix 1, strand 1, helix 4, and helix 5 (H1, S1, H4, H5), in response to ligand binding. Upon ATP binding, the lid of all hsp90s moves from the open to the closed conformation, covering the bound nucleotide and potentiating a closed dimer state. While this movement is common to all hsp90 paralogs, the movements to expose Sites 1, 2, or 3 in response to inhibitory ligands are paralog specific. Thus, for Site 1, which in Grp94 is lined by residues Leu104, Leu163, Phe199, Val209, Trp223, Ile247 (hHsp90 residues Leu48, Leu107, Phe138, Val148, Trp162, Val186), access involves the remodeling of the linker between helices 3 and 4 (H3, H4) to reposition Leu163 (Hsp90 Leu107). This conformational shift occurs in all four paralogs, but in Grp94 it results in structural disorder along H4 that lowers the affinity for moieties targeting Site 1. Site 3 is the outermost of the three pockets. The mouth of Site 3 is constricted by Thr171 and Gly196 (Thr115 and Gly135 in Hsp90). Access to Site 3 in Grp94 can be expanded by shifts in Thr171 and an unwinding of the start of H4. The equivalent motions have not been observed in Hsp90, however, and ligands that bind in Site 3 in Hsp90 exhibit energetically unfavorable conformations or interactions compared to the poses observed when binding to Site 3 in Grp94._22-24._

While ligands can access Sites 1 and 3 in all paralogs, crystal structures of Grp94 in complex with inhibitors occupying Site 2 have been instrumental to understanding how Site 2 binding favors Grp94 inhibition over Hsp90^15,19^. Ligand access to Site 2 was first identified in Grp94 ^15^ and subsequently in fungal Hsp90s ^25^. Site 2 is bordered by Leu104, Leu163, Phe199, Ala202, Phe203, Val209, Val211, Ile247, and Leu249 in Grp94, and is more interior and hydrophobic than Sites 1 or 3. In apo Grp94, Site 2 is blocked by the side chain of Phe199. Exposure of the site requires a 2.8 Å displacement of the backbone and a ∼25 degree rotation of the side chain. In the densely packed interior of the NTD, this movement precipitates a cascade of steric remodeling events that impact adjacent residues, first to Tyr200 and subsequently to H1, S1, H4, and H5 (**Figure 1B, 1C**) that leaves the lid, and particularly H1, in distinct new conformations. The human Hsp90 equivalent of Phe199 is Phe138. Phe138 has not been observed to undergo similar displacements and expose Site 2 in metazoan Hsp90, forcing ligands with a preference for Site 2 into unfavorable conformations or occupancy of alternate binding pockets such as Site 1 ^15,19^. Thus, targeting Site 2 has been significant to the development of Grp94-selective inhibitors.

The ability of the lid to move and expose Site 2 when encountering Site 2 binders is a characteristic of Grp94 that is absent in Hsp90. However, the structural origins of these differences are unknown. The lid subdomain is comprised of two discrete sections, with S1/H1 comprising the first part and H4/H5 as the second, and both of these sections are remodeled when an inhibitor binds in Site 2. Here, we ask whether we can separate the contributions of S1/H1 from the contribution of H4/H5 and what effect the uncoupling of these parts has in binding of Site 2 ligands. Our approach involved the use of inhibitors whose structures with both Hsp90 and Grp94 have been experimentally determined. We first determined the structure of PU-H36, a Site 2 directed inhibitor, bound to both Grp94 and Hsp90 as well as the structure of the high affinity pan-hsp90 inhibitor PU-H71 in complex with Grp94. We then created chimeric chaperones by swapping the S1/H1 segments of Grp94 and Hsp90 but keeping the H4/H5 of each paralog in their WT state. We tested the response of these chimeras to inhibitors that target Site 2 and Site 1. From this study, we identified H1 as the protein element that plays a critical role in determining ligand binding characteristics as well as ATPase activity between the paralogs.

## Materials and Methods

### Protein purification

Canine Grp94, which is 98% identical with human Grp94, residues 69–337 with the charged linker (residues 287–327) replaced with 4 Glycines (Grp94NΔ41) was overexpressed in *E. coli* strains BL21 Star (DE3) and purified as previously described ^22^. The purified protein was stored in 10 mM Tris, pH 7.6, 100 mM NaCl, 1 mM DTT at 30 mg/ml were stored at −80 °C. The N-terminal domain of human Hsp90α residues 1–236 was overexpressed and purified as previously described ^19^. The purified protein was concentrated to 20 mg/mL and stored at −80 °C in a buffer of 10 mm Tris, pH 7.6, 100 mM NaCl, and 1 mM DTT. Protein concentrations were determined by A280 absorption (NanoDrop 2000) and using their calculated extinction coefficients (EXPASY ProtParam).

### Ligands

PU-H36, PU-H54, and PU-H71 were synthesized as previously reported ^15,26^ and were dissolved in DMSO, typically at concentrations of 50-100 mM. bme-BnIm was a gift from Dr. Brian Blagg (University of Notre Dame). Radicicol was purchased from Sigma.

### Crystallization

Protein:ligand complexes were formed by mixing a three to five-fold molar excess of inhibitor to the concentrated protein and incubating the mixture for one hour. Initial crystallization conditions were identified using a high throughput screen at the HWI National Crystallization Center ^27^. Crystallizations were carried out using the hanging drop vapor diffusion method using equal volumes of reservoir cocktail and inhibitor-protein solution.

Crystals of Hsp90N:PU-H36 grew at 4 °C over a reservoir containing 0.1 M sodium cacodylate (pH 6.5), 0.18-0.22 M MgCl_2_, and 8-25% polyethylene glycol 2000 monomethyl ether (PEG 2K MME). Crystals typically appeared in 1-2 days and were cryoprotected by rapid sequential passage through the reservoir solution followed by the reservoir cocktail where the PEG 2K MME was increased to 35%. Cryoprotected crystals were recovered in nylon loops and immediately flash cooled in liquid nitrogen.

### Lid segment chimeras

All chimeras were generated by cross-over PCR reactions. The Grp94N^hspS1/H1^ chimera is Grp94N (69-337Δ41) with residues 69-95 replaced with Hsp90 residues 1-39. Hsp90^grpS1/H1^ chimera is Hsp90 (1-236) with residues 1-39 replaced by Grp94 residues 69-95. Grp94N^hspH4/H5^ is Grp94N (69-337Δ41) with residues 186-191 replaced with Hsp90 residues 124-130. Hsp90^grpH4/H5^ is Hsp90 (1-236) with residues 124-130 replaced by Grp94 residues 186-191. Chimeric protein expression and purification was as described above.

### Isothermal Titration Calorimetry

ITC titrations were carried out on a VP-ITC (Microcal) at 25 °C as previously described ^19^. The feedback mode was set to high and the reference power set at 10 μCal. For most of the titrations, the protein solution was loaded in the syringe and the inhibitor in the cell. Both protein and inhibitor solutions were prepared in matched buffers of 40 mM Hepes (pH 7.5), 100 mM NaCl, and 2% DMSO. Titrations involved 29 injections of 10 μL (2 μL for the first injection) set 5 minutes apart and stirring speed set at 310 rpm. The first injection was discarded in all titrations. Heat of dilution of the titrant in the syringe, referred to as background or reference titration, was subtracted two ways. If saturation was achieved, heats of dilution were estimated from the final injections, otherwise a protein into buffer background titration was performed. The concentrations of inhibitor stocks to be used for a series of titrations were determined by titrating them against known and well-behaved 1:1 binders such as Hsp90N or Grp94Nd41. Data were fit to a one-site model using Origin Software. K_d_ and other thermodynamic values were derived from the average of two replicate titrations, ± SD.

### Data collection and structure refinement

X-ray diffraction data was collected at SSRL beamline 9-2 and APS beamline 23-ID-B. Data were reduced and scaled using either HKL2000 or AutoProc 1.0.5 ^28–30^. Structures were solved by molecular replacement with either PDB 2FWZ as the search model for Hsp90N or the core region of Grp94 for Grp94N crystals. Missing residues were manually rebuilt and refined by using COOT and PHENIX ^31,32^. Inhibitor and solvent positions were identified from overlapping peaks of difference density. Ligand parameter and topology files were generated using Phenix eLBOW or the Dundee PRODRG server ^33^. Structure validation was performed with MolProbity ^34^. Data collection and refinement statistics are shown in **Table 2**. Molecular graphics were generated using PyMOL (Schrodinger).

### Thermal stability assay

A typical reaction contained 45 μL of a 5 or 10 μM protein sample mixed with 5 μL of 200X SYPRO Orange (Invitrogen). For NTD proteins the buffer contained 10 mM Tris, pH 7.6, and 100 mM NaCl. For full-length proteins the salt was increased to 150 mM. Each sample was assayed in triplicate and assays were repeated at least twice. The experiments were performed in a Stratagene MX3005P QPCR System using a stepwise temperature increase program from 25 to 99 °C (143 cycles, 0.5 °C/cycle). Melting temperatures were determined using the Boltzmann fit function in Prism 5 (GraphPad Software Inc.).

### FP Saturation Binding and Competition Assays

FP assays with the fluorescent probe Geldanamycin-FITC (Gdm-FITC, Enzo Life Sciences) were performed on a Biotek Synergy H4 hybrid microplate reader (Agilent) with black 96-well microplates (Corning 3991). Excitation and emission wavelengths were set at 485 nm and 528 nm, respectively. Assay buffer contained 20 mM HEPES-KOH, pH 7.3, 50 mM KCl, 2 mM DTT, 5 mM MgCl_2_, 20 mM NaMoO_4_, and 0.01% NP-40 with 0.1 mg/ml Bovine gamma globulin.

The saturation binding curve of Gdm-FITC with each protein sample was determined by binding 5 nM of the probe (diluted into buffer) with increasing concentrations of protein (from 0.5 nM to 256 nM) in triplicate wells. The total volume of the assay in each well was 100 μL. Plates are covered with parafilm and foil. After 20 hours at 4 °C on a shaker the resulting FP values were recorded.

We modified the FP competition experiments as follows: In duplicate wells, 80 μL protein in assay buffer is mixed with a set of serially diluted inhibitor samples (10 μL in 50% DMSO/assay buffer) in a 96 well plate and incubated for 1 h at room temperature followed by 15-20 minutes at 4 °C with gentle shaking. Gdm-FITC (10 μL of a 50 nM solution in assay buffer) is then added to the reactions. The total volume is 100 μL per well and the final protein concentration is 6 nM. Plates are covered with parafilm and foil. After incubating the plate for 20 hours at 4 °C on a shaker the FP values were recorded. IC_50_ values reported are the average from two experiments ± SD.

### ATPase assays

The relative ATPase rates of wild type and chimeric protein samples were determined with a Piper assay kit (Fisher Scientific). Protein stocks used for the assay were in 1X ATPase buffer (40 mM HEPES-KOH, pH 7.5, 150 mM KCl, and 5 mM MgCl_2_). Briefly, protein samples were mixed with increasing concentrations of ATP (Sigma) in black 96-well microplates (Corning 3991). The total volume of the protein/ATP mixture is 50 μL. An equal volume of Piper reaction mix is added to these wells before the plate was covered with parafilm and foil and incubated at 37 °C for 2 hours. The reaction was stopped by placing the plate on ice. All protein samples have a final concentration of 5 μM and ATP ranged from 12.5 to 800 μM. When all the components are added, the reaction buffer consisted of 30 mM HEPES-KOH, pH 7.5,12.5 mM Tris-HCl, pH 7.5, 112.5 mM KCl, and 3.75 mM MgCl_2_. Fluorescence intensity values were recorded on a Biotek Synergy H4 hybrid microplate reader (Agilent) with emission and excitation wavelengths set at 490 and 590 nm, respectively.

## Results

### The 8-aryl group of PU-H36 binds in Site 1 of Hsp90 and in Site 2 of Grp94

PU-H36 is a purine-based inhibitor that is selective for Grp94 ^15^. Compared to PU-H54, it incorporates a bulkier 2,4,6-trimethy 8-aryl moiety and the acetylenic pent-4-yn-1-yl tail is attached at the N9 position of the purine base instead of the N3 position found in PU-H54 (**Figure S1**) ^15^. Together these changes correlate with a ∼25-fold improvement in K_d_ for PU-H36 over PU-H54 binding to Grp94, as well as a ∼1.5-fold improvement in Grp94 selectivity over Hsp90 (**Table 1**). While modeling studies based on the structure of the Grp94 N-terminal domain (Grp94N) bound to PU-H54 are consistent with the insertion of the 8-aryl group of PU-H36 into Site 2, the paucity of structures of PU-based compounds bound to Grp94 represents a gap in our understanding of the modes of binding available to these compounds. In particular, the structural constraints imposed by the narrow Site 2 channel, combined with the bulkier 8-aryl group of PU-H36 and the significant variation in binding affinity and Grp94-selectivity for different PU compounds led us to ask how PU-H36 is accommodated in the Grp94 binding pocket and how the Grp94 conformation adapts to the different ligand. In addition, we wanted to determine the structural effect of changing the attachment site of the acetylenic tail from the purine N3 to the purine N9. To answer these questions, we co-crystallized the Grp94N:PU-H36 complex and determined its structure at 2.8 Å resolution.

**Table 1.**
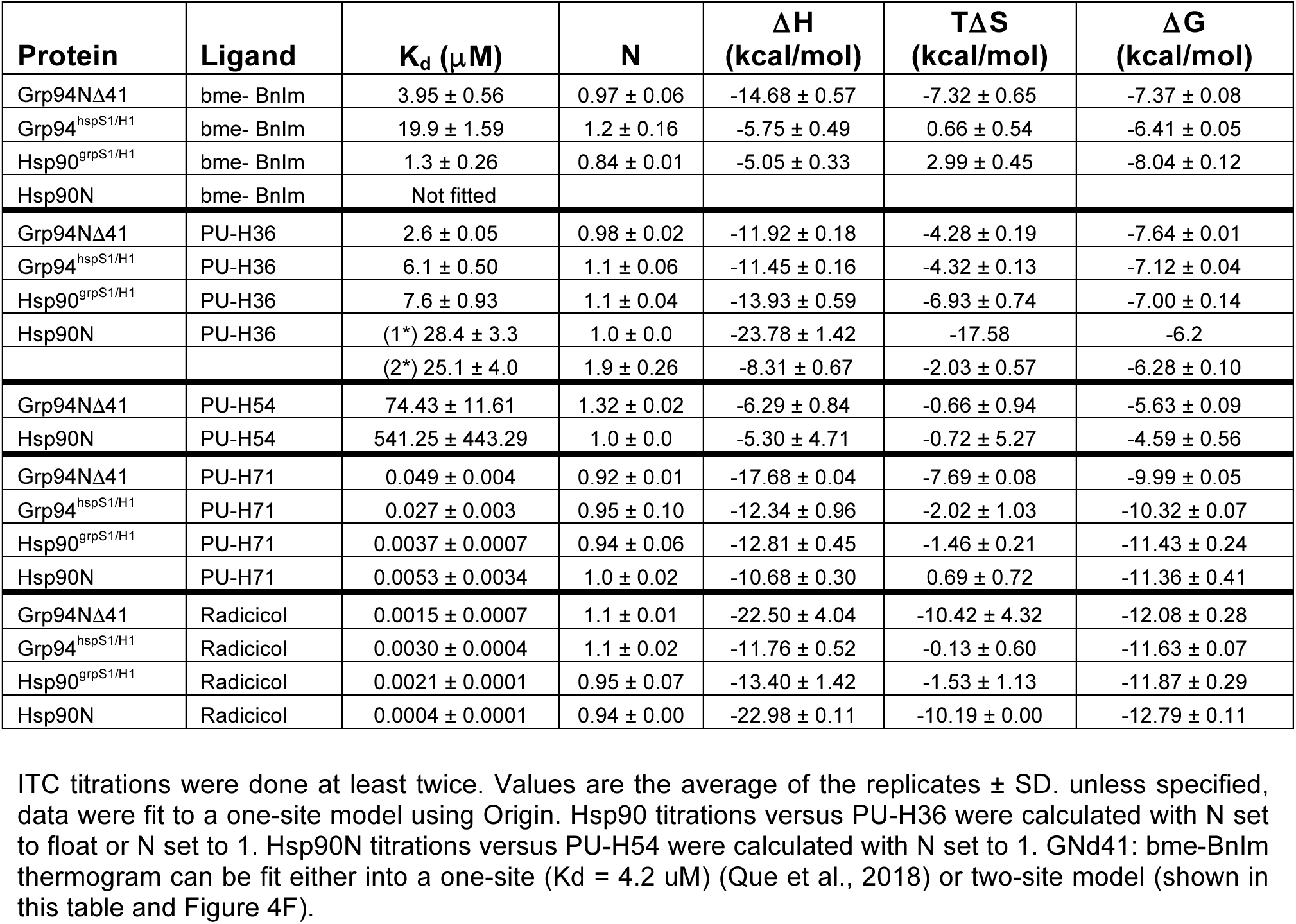
Thermodynamic parameters of binding from ITC titrations.

**Table 2.** – Data collection and Refinement Statistics.

| | Hsp90α:**PU-H36** | Grp94N:**PU-H36** | Grp94N:**PU-H71** |
| --- | --- | --- | --- |
| PDB code | 8SBT | 8TF0 | 8SSV |
| Source | SSRL 9-2 | SSRL 9-2 | APS 23ID-B |
| Space group | I 2 2 2 | P 2<sub>1</sub> 2<sub>1</sub> 2<sub>1</sub> | P 2<sub>1</sub> |
| Unit cell: a, b, c, (Å); | 66.61, 90.66, 98.48 | 96.28, 104.39, 172.28 | 51.90, 65.43, 75.38 |
| Angles ($\alpha, \beta, \gamma$) | 90, 90, 90 | 90, 90, 90 | 90, 95.20, 90 |
| Resolution (Å) | 50.00-1.50 | 50.00-2.80 | 75.07-1.72 |
| Last shell (Å) | 1.55-1.50 | 2.90-2.80 | 1.93-1.72 |
| Total (Unique) reflections | 347,644 (91,288) | 276,695 (43,634) | 125,702 (23,864) |
| Completeness (last shell) (%) | 99.9 (100.0) | 99.9 (100.0) | 91.3 (66.6) (ellipsoidal)<br>44.4 (7.5) (spherical) |
| Average I/$\sigma$ (last shell) | 48.31 (2.30) | 22.77 (1.57) | 8.9 (1.8) |
| Redundancy | 3.8 | 6.3 | 5.3 |
| R<sub>sym</sub> (last shell) | 0.040 (0.819) | 0.107 (0.841) | 0.106 (0.821) |
| R<sub>meas</sub> (last shell) |  |  | 0.118 (0.915) |
| R<sub>im</sub> (last shell) |  | 0.107 (0.841) | 0.051 (0.396) |
| CC1/2 (last shell) | n/d | n/d | 0.997 (0.655) |
| **Refinement** |  |  |  |
| Resolution range (Å) | 33.35-1.50 | 35.85- 2.80 | 75.07-1.72 |
| Reflections | 47,416 | 43,495 | 23,857 |
| Non-solvent atoms (no H) | 1,768 | 6,698 | 3,099 |
| Solvent and heteroatoms | 387 | 166 | 310 |
| RMSD Bond lengths (Å) | 0.016 | 0.002 | 0.006 |
| RMSD Bond angles (°) | 1.469 | 0.50 | 0.915 |
| R<sub>work</sub> (%) | 0.1732 | 0.228 | 0.209 |
| R<sub>free</sub> (%) | 0.1965 | 0.260 | 0.252 |
| Structure solution | MR | MR | MR |
| All atom clashscore (B < 40) | 3.34 | 6.25 | 5.01 |
| Ramachandran favored (outlier) (%) | 97.1 (0.0) | 95.1 (0.35) | 95.5 (0.0) |
| Overall score | 1.28 | 1.70 | 1.58 |

The crystallographic asymmetric unit consists of four Grp94N:PU-H36 complexes with pseudo-equivalency between the A/B and C/D pairs, yielding two independent views of the complex. As seen in **Figures 2A, 2B, 2C**, the 8-aryl group of PU-H36 is inserted into Site 2, with the plane of the aryl moiety and the 2- and 4-methyl groups closely aligned with their equivalents in PU-H54 (PDB code 3o2f). The most notable difference in the two complexes is found in the conformation of helix 1 (H1) in the lid subdomain. In the PU-H36 complex, H1 shifts from its apo protein conformation and adopts either an angled “up” or “down” trajectory. Notably, in both configurations it remains associated with the body of the N-terminal domain (NTD) (**Figures 2B, 2C**). In the PU-H54 complex, by contrast, H1 is displaced outwards and loses its association with the body of the NTD (**Figure 2D**). These differences can be ascribed to changes in the placement of Tyr200 between the two complexes. In the PU-H36 complex, Tyr200 forms edge-on-pi interactions with Phe199. In the PU-H54 complex, however, the connection between Tyr200 and Phe199 is lost. Instead, Tyr200 is rotated outwards by an additional 77 degrees, forcing the further displacement of H1 from its position in the apo form of the protein (**Figures 1B, 2E**).

**FIGURE 2.**
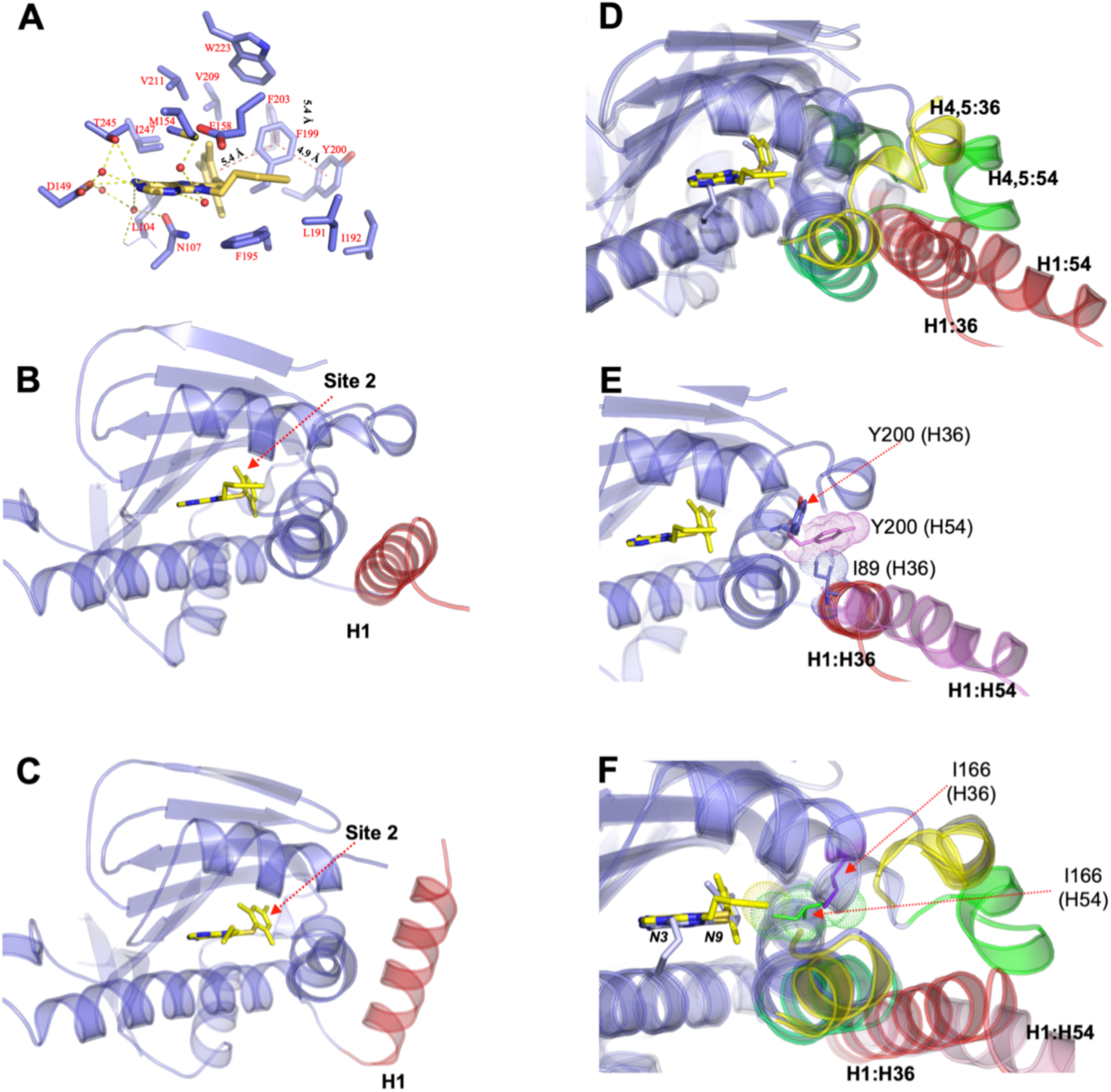
PU-H36 binding to Grp94. (A) Interactions of PU-H36 with the ATP binding pocket and Site 2. (B,C) The two unique monomers in the GrpN94:PU-H36 crystallographic asymmetric unit show the lid conformations adopted by the protein. H1 is shown in red. (D) Overlay of Grp94N:PU-H36 (slate) with Grp94N:PU-H54 (light blue). H1s (red), H4 of Grp94N:PU-H36 (yellow), H4 of Grp94N:PU-H54 (green). (E) Changes in the placement of Tyr200 between the two complexes impacts H1 conformation. (F) The attachment of the acetylinic tail of PU-H36 (yellow) at N9, in contrast to the N3-attached tail of PU-H54, contributes to the difference in H1 conformations between the two ligand bound complexes.

The reason for the differences in the placement of Tyr200 can be traced to the attachment point of the hydrophobic acetylenic tail on the two PU compounds (**Figures S1**). In PU-H36, the tail is attached to N9 instead of N3 of the purine ring. It adopts an extended conformation that projects into helix 3 (H3) while its underside is stabilized by extensive van der Waals interactions with the face of Phe195 (**Figure 2F**). This moves H3 away from the ATP binding pocket beginning at Asn162. The resulting volume created by this displacement provides space for Tyr200 to maintain its association with Phe199 after Phe199 has moved and rotated to open up Site 2. By contrast, the tail in PU-H54 is attached at the N3 position. From this more distant attachment point it cannot reach the hydrophobic interior face of H3. Instead, the N3-attached tail minimizes its solvent exposure by adopting a “scorpion tail”-like fold under the purine ring, foregoing extensive interactions with Phe195 in the process (**Figure 2F**). As a result, H3 does not move to expand the ATP binding pocket. This sterically blocks the potential Tyr200/Phe199 interaction, permitting the larger movement of Tyr200. Thus, the larger repositioning of H1 in the PU-H54 complex is ultimately tied to the conformation of the acetylenic tail, which in turn is dictated by its site of attachment on the purine ring.

The structure of Grp94N:PU-H36 also helps explain the improved binding of PU-H36 to Grp94 compared to PU-H54. A characteristic feature of all Grp94 complexes that expose Site 2 is the large rotation of Phe195 from its position sandwiched between helices 4 and 5 (H4, H5) in the apo protein, into a position that shields the underside of the ATP binding pocket from solvent. The additional interactions between the PU-H36 tail and Phe195 suggest that these contribute positively to the improved binding of PU-H36 (K_d_ =2.6 μM) for Grp94 compared to PU-H54 (K_d_ = 74 μM) (**Table 1**), which lacks these interactions. The attachment of the alkyl tail at N9 is thus more optimal for Grp94 binding as opposed to an attachment at N3.

### Binding of PU-H36 to Hsp90 compared to other PU-inhibitors

Compared to PU-H54, PU-H36 binds with higher affinity to both Grp94 and to Hsp90 and also exhibits greater fold-selectivity for Grp94 over Hsp90 (**Figure 4**, **Table 1**). In Grp94, the 8-aryl group of PU-H36 inserts into Site 2 of the ATP binding pocket, and the N3-tail makes extensive interactions with Phe195 of H5. To visualize the placement of these elements when PU-H36 is bound to Hsp90, we crystallized the human Hsp90 N-terminal domain (Hsp90N) in complex with PU-H36 and solved the structure at 1.50 Å resolution. As seen in **Figure S2A**, the protein structure is identical to Hsp90N:PU-H54 (PDB Code 3o01), with an RMSD of 0.088 Å for CA atoms. As with Hsp90-bound PU-H54, the 8-aryl group of PU-H36 is located in Site 1 of the binding pocket, rather than in the hydrophobic Site 2 as in Grp94. This likely accounts for much of the energetic penalty these compounds incur in binding to Hsp90. In addition, the acetylenic tail, which is attached at the N9 of the purine ring in PU-H36, curves under the purine moiety in the “scorpion tail” conformation in a similar manner as the identical N3-attached moiety in PU-H54 and makes no stabilizing interactions with the Hsp90 protein (**Figure S2B**). However, because the N9-attached tail of PU-H36 is more interior to the ATP binding pocket compared to the N3-attached tail of PU-H54, the displacement and disorganization of the nearby solvent molecules is less than that seen in Hsp90N:PU-H54. This may explain the slightly better binding of PU-H36 to Hsp90 than PU-H54.

Many purine-based inhibitors are not paralog selective, and unlike the Grp94-selective PU-H36, the pan-hsp90 inhibitor PU-H71 (**Figure 4G**) binds with nanomolar affinity to all hsp90 paralogs ^23,35^. A notable difference between PU-H71 and the Grp94-selective PU compounds is the larger 8-aryl group, which contains both an iodine at the 2’ position along with a 4’5’-methylenedioxy cap. While both PU-H71 and PU-H36 are Hsp90 Site 1 binders, PU-H71 binds to Hsp90 with a 5×10^3^-fold higher affinity than PU-H36 (**Table 1)**. Comparing the structures of PU-H36 and PU-H71 in complex with Hsp90, it is apparent that PU-H71 forms a more extensive set of protein-ligand interactions within the Hsp90 ATP binding pocket that provides a rationale for the observed difference in binding (**Figure 3A**). In particular, the bicyclic 8-aryl of PU-H71 allows it to make extensive pi-pi contacts with Phe138 and with both rings of Trp162. In addition, the 2’-iodine forms a water mediated hydrogen bond with the Leu107 backbone oxygen ^35^. These stabilizing interactions are not made by PU-H36 since its 8-aryl moiety is both smaller and does not have any polar or charged functionality. The curled tail conformation of PU-H36 when bound to Hsp90 (**Figure S2B**) also differs from the extended trimethylamine tail of PU-H71 (**Figure 3A**). The charged tail of PU-H71 not only increases the ligand’s solubility, but it also allows for additional hydrogen bond interactions in the pocket. The hydrophobic alkyl tail and 8-aryl moiety of PU-H36 cannot duplicate these stabilizing interactions shown by PU-H71.

**FIGURE 3.**
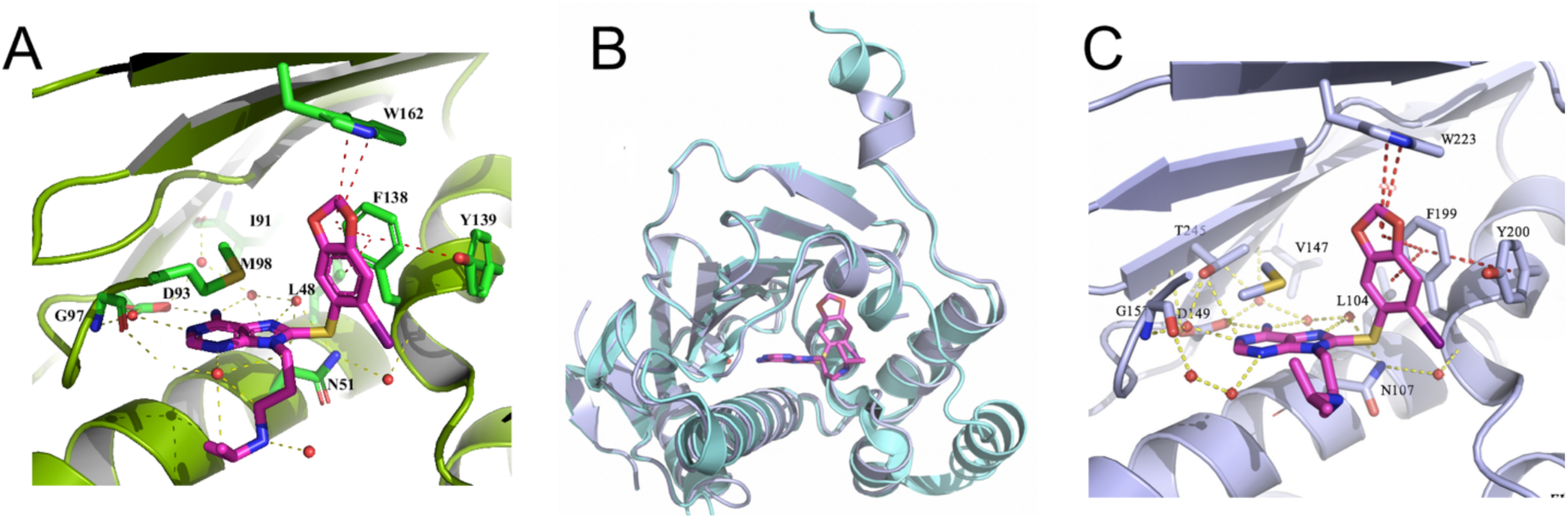
Grp94 binds PU-H71 in Site 1. (A) Interactions of PU-H71 with the Hsp90 ATP binding pocket. (B) The structure of Grp94N:PU-H71 (dark blue) is similar to apo Grp94N (cyan). While helix 3 and strand 1 of the lid was observed and can be superposed to that of the unliganded structure, portions of H4 and H5 in Grp94N:PU-H71 are disordered and are shown as dash lines. (C) PU-H71 in the Grp94 main pocket is stabilized by a network of hydrogen bonds (yellow dashes), van der Waals and pi-pi interactions (red dashes). Phe199 is in the same position as in apo Grp94 and the 8-aryl of PU-H71 is found in Site 1.

**FIGURE 4.**
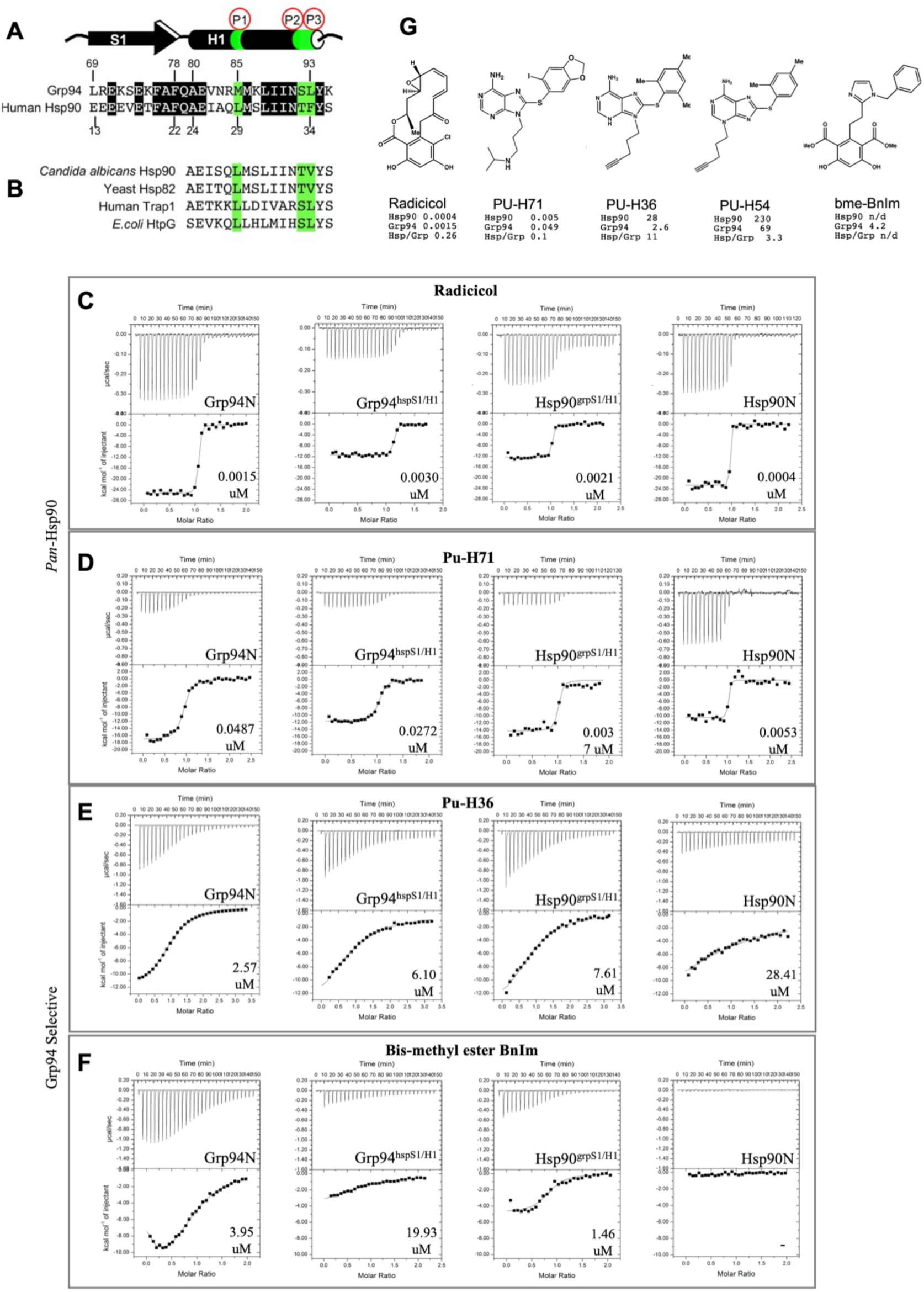
The effect of S1/H1 swaps on ligand binding. (A) Comparison of S1/H1 between Grp94 and Hsp90. Identical residues are shaded black, key discriminating residues in H1 are shaded green. (B) H1 residues for selected Hsp90 orthologs and paralogs. (C-F) Representative ITC thermograms for Grp94, Hsp90, and the S1/H1 chimeras titrated with different inhibitors. Titration with pan-inhibitors radicicol and PU-H71 (C,D) and with Grp94 selective ligands PU-H36 and bme-BnIm (E,F). Kd values are the average of two replicate titrations, ± SD. See **Table 1**. (G) Ligands discussed in this report.

### Structure of Grp94N with PU-H71 reveals Site 1 binding

Unlike PU-H36 and PU-H54, which bind with higher affinity to Grp94 than to Hsp90, PU-H71 exhibits a ∼9-fold greater affinity for Hsp90 over Grp94 (**Table 1**) ^23,35^. In Hsp90 the 8-aryl group is inserted into Site 1 and derives its high affinity from extensive contacts in that pocket. For Grp94, modeling studies show that the 8-aryl group of PU-H71 could be accommodated into either Site 1 or Site 2 upon the previously described remodeling of the ATP binding pocket. In order to understand how PU-H71 binds to Grp94, and to explore the basis for its lower affinity compared to Hsp90, we determined the co-crystal structure of PU-H71 in complex with Grp94N at 1.72 Å resolution.

The structure of Grp94N:PU-H71 (**Figure 3B**) reveals that the 8-aryl group of PU-H71 is inserted into Site 1 of the Grp94 ATP binding pocket. Site 2 remains closed, and without the extensive rearrangements needed to accommodate the movement of Phe199, Tyr200, and H1, the overall protein conformation is similar to that of apo-Grp94 (PDB code 1yt1). A small unwinding at the end of H3 involving residues 161-163 occurs as a consequence of accommodating the larger 8-aryl moiety in Site 1. Density corresponding to residues 164-187 of H4 was observed during refinement but the quality was insufficient to include in the final model.

When bound to Grp94, PU-H71 adopts a pose similar to that found in Hsp90 (**Figure 3A, 3C**). An identical architecture of stabilizing interactions found in Hsp90N:PU-H71 is also observed in Grp94N:PU-H71, including hydrogen-bonding of the purine N6 with Asp149, water-mediated H-bonds to Leu104, Asn107, Val147, Asp149, Gly153, and Thr245, and the hydrophobic contacts between Met154 as well as Ala111 and the adenine ring. Similar to that found in Hsp90N:PU-H71, the ligand displays an extensive network of pi-pi interactions with Phe199, Tyr200, and Tryp223. Thus, the same factors that allow PU-H71 to achieve nanomolar binding to Hsp90 are also at work in binding to Grp94. However, the partial disorder of H4 of Grp94, as was noted previously for the compound SNX-0723 ^36^, likely exacts an energetic penalty by weakening the interactions in the adjacent Site 1, leading to the observed decrease in binding relative to Hsp90 (**Figure 4D**, **Table 1**).

### Effect of swapping strand 1 and helix 1 on Site 2 ligand binding

Based on the available structures of Grp94 and Hsp90 bound to selective and non-selective ligands, the “lid” subdomain can be regarded as having two discrete sections, with S1/H1 comprising the first part and H4/H5 as the second. While only H4/H5 is observed to rearrange when a ligand is inserted in either Site 1 or Site 3, the entire lid subdomain, including S1/H1, is remodeled when an inhibitor utilizes Site 2.

A sterics-based mechanistic explanation for this observation comes from an analysis of Grp94 structures co-crystallized with Site 2 ligands (PU-H54, PU-H36, BnIm, Bis-methylester-BnIm, Bis-chloro-BnIm) ^15,19^. These structures reveal that access to Site 2 depends on the backbone of Phe199 pivoting 2.8 Å away from its apo position and the side chain rotating ∼25 degrees to allow for a hydrophobic moiety to be inserted into this exposed pocket. The movement of Phe199 forces Tyr200 to shift in order to prevent steric clashes, ultimately triggering a cascade of conformational rearrangements involving S1/H1 and the neighboring helices H4 and H5, resulting in Grp94 lid conformations that differ significantly from those found in the apo or ATP-bound protein. In Hsp90, a similar flexibility of S1/H1 has not been observed, and the Grp94 Site 2-targeted moieties of ligands, such as the 8-aryl groups of PU-H54 and PU-H36, are instead confined to Site 1 in this paralog.

Because only Grp94, and not Hsp90, is capable of the wholesale remodeling of the lid that is associated with Site 2 binding, we asked whether we can separate the contribution of S1/H1 to this binding from that of H4/H5. In particular, we asked whether differences in S1/H1 alone were sufficient to change the mobility of the lid and thus allow access to Site 2 (**Figure 1B**). The S1/H1 segment consists of 27 amino acids, of which 13 differ between Hsp90 and Grp94 (**Figure 4A**). To test this hypothesis, we created chimeric proteins, termed Hsp90^grpS1/H1^ and Grp94^hspS1/H1^, where S1/H1 from one paralog was substituted with S1/H1 from the other while keeping the H4/H5 of each paralog in their WT state. We then used ITC to determine the effect of the swap on the binding of inhibitors that preferred or required access to Site 2.

We tested two different Site 2 inhibitors: 1) PU-H36, which exhibits 10-fold selectivity for Grp94 when its 8-aryl group is inserted into Site 2, compared to its binding to Site 1 in Hsp90; and 2) bis-methylester BnIm (bme-BnIm), an inhibitor whose 1- and 3-methyl ester moieties on the resorcinylic scaffold forces it to interact with Site 2 and, more importantly, exhibits no detectable binding to Hsp90 by ITC (**Figure 4E, 4F, Table 1**). As controls, we tested radicicol, a high affinity inhibitor that binds in the ATP pocket but does not utilize either Site 1 or Site 2, and PU-H71, a high affinity, non-selective inhibitor that we have shown above binds in Site 1 of both Grp94 and Hsp90 (**Figure 4C, 4D, Table 1)**. With the structural data presented in this report, all four ligands have crystal structures in complex with Grp94 and with Hsp90 (Grp94N:bme-BnIm PDB code 6baw; Hsp90N:bme-BnIm PDB code 6ceo; Grp94N:Radicicol PDB code 1u0z; Hsp90N:Radicicol PDB code 4egk; Hsp90N:PU-H71 PDB code 2fwz) ^19,22,35,37^, allowing us to correlate S1/H1 and ATP pocket remodeling with binding affinity.

PU-H36 binds to Grp94 with a K_d_ of 2.6 μM and to Hsp90 ∼10 fold less tightly (**Figure E, Table 1**). The Hsp90N:PU-H36 structure shows that the 8-aryl moiety of the ligand is in Site 1 and that the lid, including S1/H1, is unchanged from WT. When S1/H1 is replaced with the equivalent element from Grp94 (Hsp90^grpS1/H1^), however, PU-H36 binding to the chimera improves from a K_d_ of 25-28 μM in WT Hsp90 to a K_d_ of 7.6 μM, corresponding to a free energy change of −0.72 kcal/mol.

The observed gain in affinity for PU-H36 by Hsp90^grpS1/H1^ suggests that the S1/H1 swap has caused this chimera to acquire a different mode of binding for PU-H36. In particular, the observed increase in affinity is consistent with the conformational flexibility of S1/H1 from Grp94 being imparted to Hsp90^grpS1/H1^, thereby allowing the 8-aryl group of PU-H36 to access Site 2 of the chimera. This interpretation follows from considering two alternative scenarios. In the first alternate scenario, the 8-aryl moiety of PU-H36 sits in Site 1 of Hsp90^grpS1/H1^, recapitulating that seen in WT Hsp90. The impact of the S1/H1 swap on binding and its resulting affinity with PU-H36 is predicted to be minimal, however, in contradiction to what is observed, since binding to Site 1 only involves interactions with H4/H5, which are unchanged in the Hsp90^grpS1/H1^ chimera. Alternatively, if the S1/H1 in this chimera is more flexible but does not result in opening of Site 2, then the entropic penalty of a floppy S1/H1 should lead to lower, not higher, affinity. Together, these observations suggest that the structure of this chimera now allows access to Site 2. The deduced interaction with Site 2 in the chimera is also consistent with earlier studies with PU derived compounds that showed that while the purine positions the ligand in the main ATP pocket, other portions of the PU analogs, particularly those interacting with the lid, are the key determinants of binding affinity to Hsp90 ^38^.

In contrast, the corresponding Grp94 chimera, Grp94 with S1/H1 from Hsp90 (Grp94^hspS1/H1^) exhibits a K_d_ of 6.1 μM for PU-H36, which is a 2.4-fold loss in affinity relative to WT Grp94. The corresponding free energy change is +0.52 kcal/mol, which is of a similar magnitude but opposite sign of the S1/H1 swap into Hsp90 discussed above. This suggests that an alteration in how the 8-aryl moiety is accommodated in the cavity of Grp94^hspS1/H1^ has occurred. If the 8-aryl moiety of PU-H36 has transitioned into Site 1 of chimeric Grp94^hspS1/H1^, this change from the hydrophobic Site 2 pocket to a more solvent accessible environment in Site 1 would extract an energetic penalty that could explain the loss of affinity compared to WT. PU-H36 with its 8-aryl group in Site 1 of Grp94 can be modeled into the Grp94N:PU-H71 structure, where PU-H71 is in Site 1 (**Figure S4**) and S1/H1 is not changed from its apo conformation.

To further investigate the effects of the H1/S1 swap on Site 2 accessibility, we tested bme-BnIm, a resorcinylic inhibitor that triggers Site 2 pocket opening and lid remodeling upon binding to Grp94 (**Figure S3**) ^19^. Compared to PU-H36, where the flexible 8-aryl group can adopt a high affinity ‘backward-bent’ pose by inserting into Site 2, or a lower affinity “forward-bent” pose by accessing Site 1 (**Figure S1**), the bulky methyl esters attached to the resorcinylic framework of bme-BnIm are fixed in their configurations and prevent the adoption of an alternate pose (**Figure S5A**). In Grp94, these fixed configurations place the 1-methyl ester in Site 2 and the benzyl imidazole in Site 1. While bme-BnIm binds to Grp94 with a K_d_ of 3.95 μM, ITC experiments showed a nearly complete loss of enthalpy and affinity for Hsp90 (**Figure 4F**). Confirming this loss of binding, the structure of bme-BnIm soaked into crystals of Hsp90 showed that the resorcinylic scaffold is displaced in the binding pocket and the pendant benzyl imidazole is disordered, all of which are hallmarks of non-specific, low-affinity binding (**Figure S5B**) ^19^. Thus, because both structure and binding assays show that bme-BnIm has no low-affinity alternate pose available for productive binding in Hsp90, this ligand can be used to probe the effect of a S1/H1 swap on the plasticity of the binding pocket, particularly Site 2, in the Hsp90^grpS1/H1^ and Grp94^hspS1/H1^ chimeras.

Swapping S1/H1 of Grp94 into Hsp90 shows a remarkable change in its binding of bme-BnIm. The ITC thermograms of WT Hsp90 compared to chimeric Hsp90^grpS1/H1^ with bme-BnIm show a change from a flat binding curve with very small enthalpy to a sigmoidal one representing a binding event (**Figure 4F**). A large gain in binding affinity for the ligand by the chimeric Hsp90^grpS1/H1^ is also noted when compared to WT Hsp90 (**Table 1**). Because the resorcinylic scaffold can only make productive interactions with the binding pocket when the 1-methyl-ester of bme-BnIm inserts into Site 2, this change in affinity suggests that Site 2 in chimeric Hsp90^grpS1/H1^ has opened up as a consequence of the S1/H1 swap. Interestingly, the K_d_ of chimeric Hsp90^grpS1/H1^ for bme-BnIm (1.3 μM) is slightly better than that of WT Grp94 (3.95 μM), possibly reflecting the improved stabilization of the benzyl imidazole in Site 1 of Hsp90^grpS1/H1^ compared to WT Grp94, where part of H4 is disordered ^19^.

The opposite effect is observed when the Hsp90-derived S1/H1 is substituted into Grp94. Compared to the increase in affinity seen in Hsp90^grpS1/H1^, a ∼5-fold decrease in affinity, from 3.95 to 19.9 μM (ΔΔG = +0.92 kcal/mol) for bme-BnIm is noted with Grp94^hspS1/H1^. The sigmoidal curve seen in WT Grp94 binding to this ligand is replaced by a more flattened curve. The apparent enthalpy is also much less than that with WT Grp94 (**Figure 4F**, **Table 1**). This suggests that, compared to WT Grp94, chimeric Grp94^hspS1/H1^ has a modified binding pocket, most likely involving a loss of access to Site 2 that reduces the affinity for the inhibitor. This outcome can be directly ascribed to the swapped S1/H1. The residual binding activity detected suggests that a non-optimal accommodation of the resorcinylic scaffold in the cavity is occurring, such as that shown in the Hsp90:bme-BnIm structure ^19^.

### Effect of swapping S1/H1 on the main ATP cavity and on Site 1 inhibitor binding

In control experiments, both Grp94^hspS1/H1^ and Hsp90^grpS1/H1^ displayed a small loss in binding for radicicol but retained single nanomolar affinity (**Figure 4C**, **Table 1**). This indicates a minor role for S1/H1 in situating radicicol within the ATP binding pocket, which agrees with the structures of the Grp94N:radicicol and Hsp90N:radicicol complexes (PDB codes 1u0z, 4egk) ^22,37^. In addition, these results show that the main cavity is undisturbed by the S1/H1 swap. The steep titration curves for radicicol binding seen in **Figure 4C** highlight the difficulty in accurately determining the K_d_ of ligands binding to protein with single nanomolar or tighter affinity by ITC ^39,40^ and as such the slight differences in K_d_ observed should only be interpreted conservatively as indicative that the main ATP cavity where the purine ring binds is unaffected by the swaps.

Similarly, the two chimeras exhibit only minor changes in their binding to the high affinity non-selective inhibitor PU-H71. The measured K_d_ of PU-H71 to WT Hsp90 is 5.3 nM while its K_d_ to chimeric Hsp90^grpS1/H1^ is 3.7 nM (ΔΔG = −0.21 kcal/mol). Meanwhile chimeric Grp94^hspS1/H1^ showed a similar small change in K_d_ (27.2 nM) relative to that of WT Grp94 (48.7 nM) (ΔΔG = −0.35 kcal/mol). The small difference in K_d_ between these chimeras and the WT proteins is not surprising because, as a Site 1 binder, PU-H71 does not precipitate structural changes involving S1/H1 in either Hsp90 or Grp94. In particular, PU-H71 binding does not perturb the position of the Hsp90 Phe138 (Grp94 Phe199) side chain relative to the unbound protein. The crystal structure of Grp94 with PU-H71 described above helps clarify the ITC data obtained with chimeric Grp94^hspS1/H1^ by showing that PU-H71 binding does not change the conformation of H1 relative to that in apo Grp94 (**Figure 4D**). Thus, both the ITC and structural data reveal that S1/H1 does not play a major role in stabilizing and binding of Site 1 inhibitors.

### FP competition assays confirm the effect of S1/H1 swap on inhibitor binding of Hsp90^grpS1/H1^

ITC titrations are carried out with equilibration times on the order of minutes, and may be inaccurate if the protein conformational changes required to open up Site 2 require more time to achieve equilibrium. Furthermore, ITC does not report on the site of binding, and it is possible that the low enthalpy observed when bme-BnIm is titrated into Hsp90 reflects an entropically driven event. In order to verify the gain in affinity obtained from ITC titrations as well as to confirm that the main ATP pocket is the site of binding in Hsp90^grpS1/H1^, we carried out fluorescence polarization (FP) competition assays between the Site 2 inhibitors using FITC-geldanamycin, a high affinity pan-Hsp90 inhibitor, as the displaced tracer.

The FP competition assays support the role played by S1/H1 in binding of Site 2. As shown in **Figure 5**, PU-H36 is selective for Grp94 and has an IC_50_ of 3.0 μM for Grp94 and >50 μM for Hsp90. Recapitulating the gain in affinity for PU-H36 observed in ITC titration experiments, the chimeric Hsp90^grpS1/H1^ shows a ≥5-fold improvement in binding of PU-H36 with an IC_50_ of 10.8 μM compared to wild type Hsp90 (≥50 μM).

**FIGURE 5.**
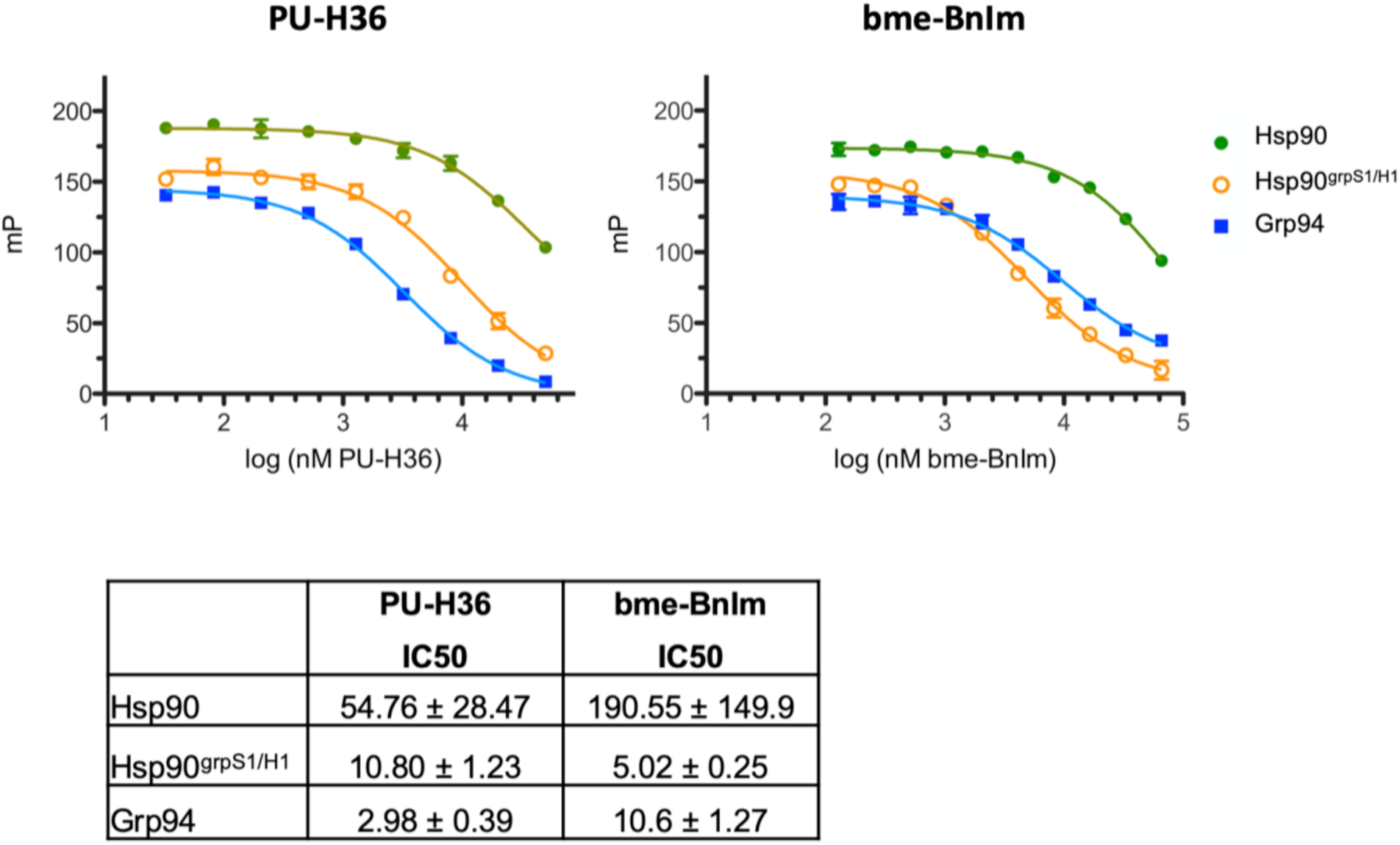
Binding of Site 2 inhibitors by the chimeric Hsp90^grpS1/H1^. FP competition assay between geldanamycin-FITC and Site 2 inhibitor PU-H36 or bme-BnIm. Representative curves are shown. IC50 values are the average of two separate replicates, ± SD. Proteins are full length or near full length constructs.

A similar trend is observed with bme-BnIm (**Figure 5**). This ligand shows a preference for Grp94 over Hsp90, with IC_50_s of 10.6 μM and >100 μM, respectively. Our results show that the S1/H1 swap in Hsp90 results in Hsp90^grpS1/H1^ having an IC_50_ of 5.0 μM, which is a ≥20-fold increase in binding affinity for bme-BnIm compared to that with WT Hsp90. Interestingly, Hsp90^grpS1/H1^ shows a slightly better IC_50_ (5.0 μM) for bme-BnIm relative to that of WT Grp94 (10.6 μM). The K_d_ values from the ITC assays also show the same slight increase in affinity for bme-BnIm by Hsp90^grpS1/H1^ compared to WT Grp94 (1.3 μM vs. 3.95 μM, respectively). Thus, data from the FP competition assays are consistent with S1/H1 being a key element in determining the plasticity of the hsp90 binding pocket, a necessary structural accommodation that leads to discriminate binding of Site 2 inhibitors.

Significantly, the impact of swapping S1/H1 between Grp94 and Hsp90 is independent of whether the isolated NTD or the full-length protein is assayed. The results from the ITC titrations, which used the NTDs of the chimeric proteins, or the FP competition assay, which use the full-length forms, are in agreement and lend further credence to the proposed role of S1/H1.

### Effect of the S1/H1 swap on thermal stability and ATPase activity

S1/H1 conformational shifts are required for the exposure of Site 2. Because conformational plasticity is a requirement for these rearrangements, we asked whether the swapping of this fragment is reflected in the overall stability of the protein. To do this, we tested the stability of the NTDs of Grp94 and Hsp90, along with the two S1/H1 chimeras, using a Thermal Shift Assay. As seen in **Figure 6A**, WT Grp94 has a T_m_ of 37.2 °C, which is lower than that of WT Hsp90 (38.9 °C). The Hsp90^grpS1/H1^ chimera, on the other hand, has a T_m_ of 31.8 °C, a 7.1 °C drop from WT that is indicative of a significant destabilization. In contrast, chimeric Grp94^hspS1/H1^ shows a small increase in T_m_ (0.8 °C) relative to that of WT Grp94, bringing it nearly on par with WT Hsp90. Together these results show that S1/H1 from Grp94 lowers the thermal stability of the protein, while S1/H1 from Hsp90 provides increased stabilization, and correlates with the ability of Grp94 S1/H1 to rearrange and allow exposure of Site 2.

**FIGURE 6.**
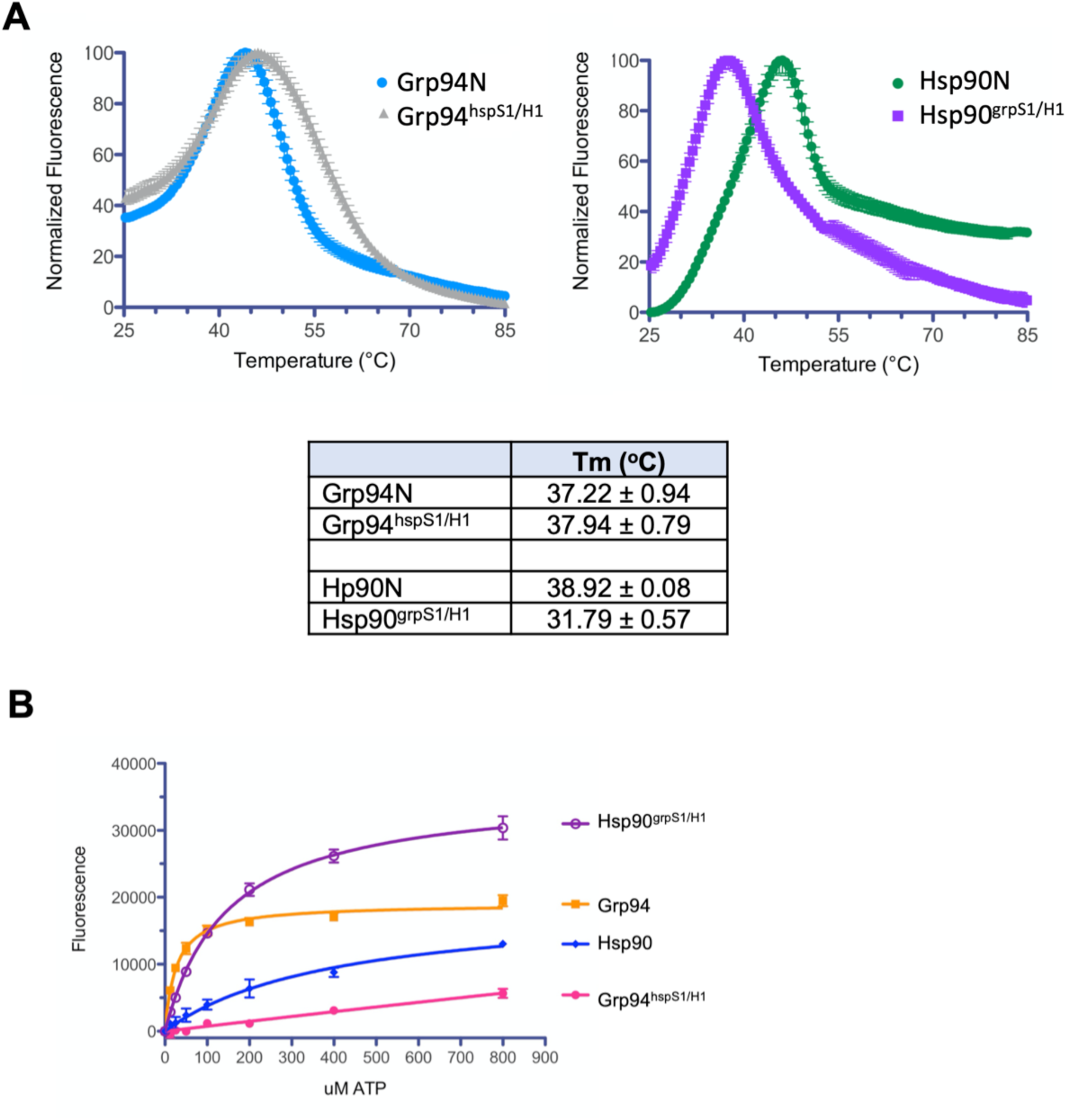
Effect of S1/H1 swaps on the thermal stability and ATPase activity of the chimeras relative to WT protein. (A) Destabilization of the Hsp90N structure occurs when S1/H1 is from Grp94. Representative melting curves of the Thermofluor assay is shown. Tm values are the average of two replicates, ± SD. (B) The chimera Hsp90^grpS^^1/H1^ retains and improves on the ATPase activity of WT Hsp90. The curves shown are from the combined fluorescence values of at least two independent replicates ± SEM.

Because of the significant decrease in the thermal stability of the chimeric Hsp90^grpS1/H1^ compared to WT Hsp90, we asked whether these S1/H1 swap chimeras retained ATPase activity. ATP hydrolysis requires the formation of the closed dimer state, where the NTDs of the two protomers are held together by cross-protomer rearrangements of S1/H1 that serve as “straps” to stabilize the closed state. Binding of ATP by all hsp90 chaperones is fast, and the rate of ATP hydrolysis is governed by the speed at which the NTDs from each protomer associate to form the closed dimer ^41,42^.

The impact of the S1/H1 swap on the ATPase activities of these chimeras is shown in **Figure 6B**. Compared to WT Grp94, the ATPase activity of chimeric Grp94^hspS1/H1^ is diminished to only ∼30% that of WT protein. The Hsp90^grpS1/H1^ chimera, on the other hand, shows a 233% increase in activity relative to WT Hsp90, and interestingly, is 150% faster than WT Grp94. Both of these altered ATPase activities can be ascribed to the swapped S1/H1, where the S1/H1 of Grp94 increases the ATPase rate while S1/H1 of Hsp90 has the opposite effect.

It is significant that these chimeras have retained the capacity to hydrolyze ATP because it demonstrates that the swapped S1/H1 of the protein does not prevent the conformational changes needed for hydrolysis. Thus, Hsp90^grpS1/H1^ allows the formation of the closed dimer and also maintains the fidelity of interactions between the N and M domains for ATPase activity.

Our results here confirm the impact of S1/H1 in determining the properties of hsp90 chaperones, including ligand selectivity and ATPase activity. The effect of S1/H1 on Hsp90 is particularly striking and suggests a notable alteration of its rigid conformation. By exchanging its S1/H1 with that from Grp94, the resulting Hsp90^grpS1/H1^ chimera has acquired ligand binding characteristics and ATPase activity that are more similar to that of Grp94 than to WT Hsp90. Thus, our data suggests that S1/H1 is the critical structural element that differentiates the characteristics between these two paralogs.

### Effect of swapping H4/H5 of the lid on inhibitor binding

Compared to Hsp90, the H4/H5 lid domain of Grp94 contains a five amino acid insertion that lengthens H4 and alters the orientation of the lid relative to the body of the NTD (**Figure S6**) ^22^. To assess the contribution of H4/H5 on selective ligand binding, and to compare it with the effects of S1/H1, we swapped H4/H5 between Hsp90 and Grp94 and tested the ability of these chimeras to bind bme-BnIm. As seen in **Figure 7**, chimeric Grp94^hspH4/H5^ binds bme-BnIm with a K_d_ of 7.5 μM, which is slightly lower than the affinity for WT Grp94 (3.95 μM). Because bme-BnIm cannot make productive interactions in the ATP pocket without access to Site 2, this similar affinity suggests that Site 2 binding in the chimera is retained. The difference in affinity (ΔΔG = 0.37 kcal/mol) may derive from the effect of the shorter H4/H5 on the stabilization of the benzyl imidazole. When comparing the effect of swapping S1/H1 and H4/H5 on bme-BnIm binding, it is clear that the effect of the S1/H1 swap (ΔΔG = 0.96 kcal/mol) is more pronounced than that due to the H4/H5 swap (ΔΔG = 0.37 kcal/mol) in binding to Site 2 binding and supports a conclusion that between the two lid segments, S1/H1 exert a bigger influence in allowing Site 2 binding.

**FIGURE 7.**
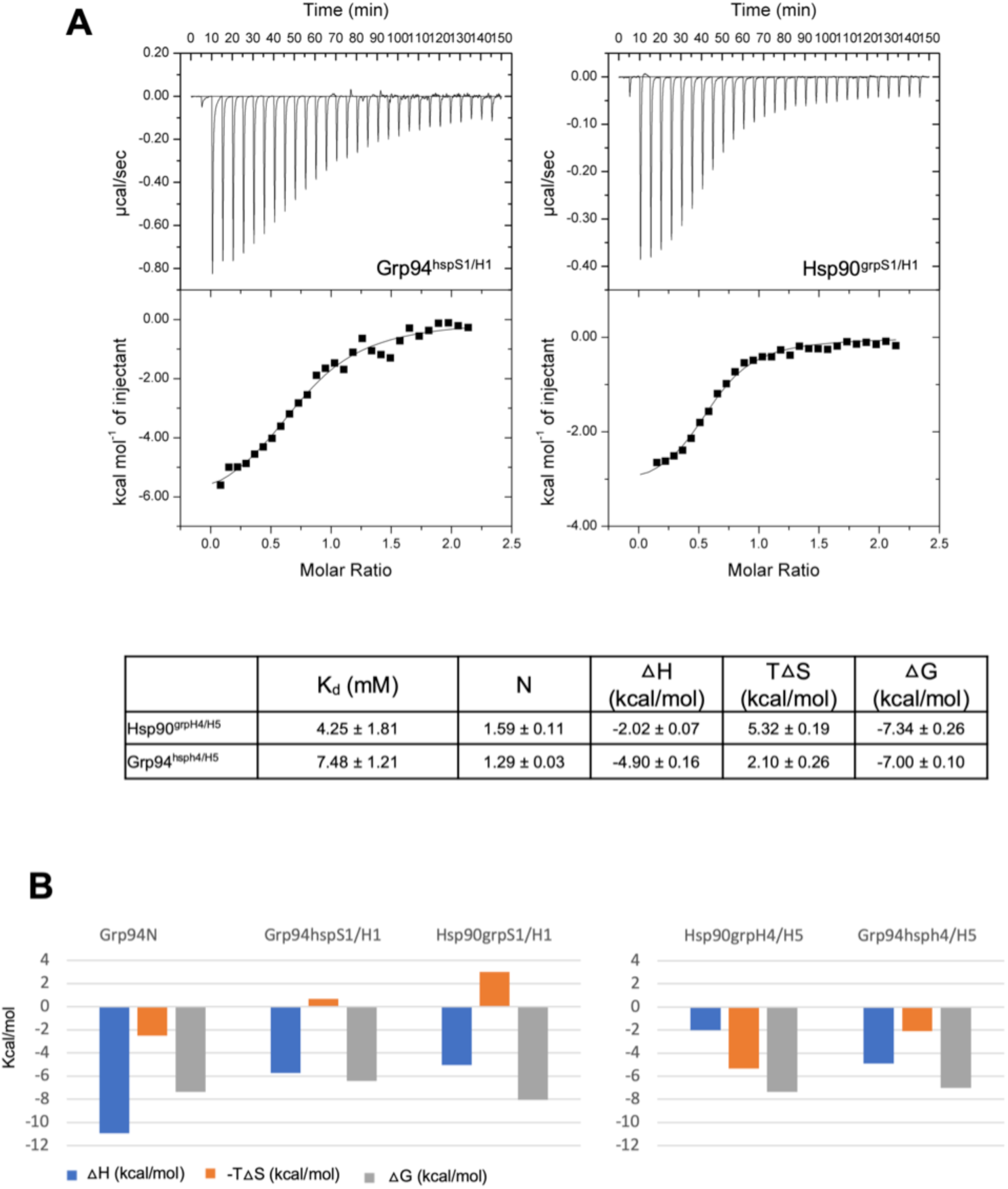
Effect of H4/H5 swap. (A) Representative ITC titrations of bme-BnIm vs. chimeras with H4/H5 of the lid swapped. Protein in the syringe is titrated into the cell containing bme-BnIm and data was analyzed as with “protein in syringe”, which results in an N value = 1/N when the protein is in the cell (normal titration). When stoichiometry is 1:1, the N should be nearly equal to 1 when calculated either way. The values in the table are the average values from two titrations of each protein sample. (B) Thermodynamics of bme-BnIm binding of the various constructs.

It was not possible to accurately determine the effect of incorporating H4/H5 from Grp94 into Hsp90. Hsp90^grpH4/H5^ was only marginally stable and precipitated during purification, upon concentration for ITC, and following recovery from ITC titration experiments.

Together, these results suggest that H4/H5 plays only a minor role in binding Site 2 targeted inhibitors and support the identification of S1/H1 as the major determinant in a paralog’s response to Site 2 ligand binding in the ATP pocket.

## Discussion

The design of paralog-selective inhibitors for the hsp90 family has been a challenge due to the high degree of amino acid identity within the ATP binding cavity. Along with the identification of conditionally accessible side pockets in this cavity came the recognition that the exposure of these regions could change the targetable landscape of this cavity. Mechanistically, however, the question of how the ATP binding cavity becomes remodeled to expose the three side pockets, and which protein structural elements are responsible for allowing the remodeling movements to take place, had remained unaddressed. While lid subdomain is the mobile element of the N-terminal domain and the most likely candidate for allowing remodeling of the ATP cavity, it has not been clear which part of the lid controls cavity remodeling. Here, we have used S1/H1 swap chimeras to uncouple the contributions of the S1/H1 and H4/H5 elements of the lid to the exposure of Site 2. We have shown that S1/H1, and not H4/H5, play the key role in determining the selective ligand binding characteristics as well as ATPase activity associated with each of the two paralogs Grp94 and Hsp90. This is particularly true of Hsp90 and our results allow us to ascribe the rigidity of the Hsp90 binding pocket that disallows binding into Site 2 as originating from the S1/H1 of the lid subdomain. This was made the evident when S1/H1 of the conformationally flexible Grp94 is placed on Hsp90. With the conformationally flexible S1/H1 from Grp94, the S1/H1 swap chimera Hsp90^grpS1/H1^ showed a remarkable gain of affinity for bme-BnIm, the Site 2 targeted inhibitor that cannot assume an alternate low energy pose in the ATP binding pocket. Our results strongly suggest S1/H1 of the lid are the major determinants of whether the NTD ATP binding pocket can undergo the conformational shifts needed to expose Site 2 upon ligand binding.

The results presented here contrast with earlier suggestions that the difference allowing the exposure of Site 2 in Grp94 resided in the five amino acid insertion that extends H4 in Grp94, an insertion that is not found in Hsp90 ^15,17,22^. Our report, together with our previous structures, suggests that H4/H5 plays a role in augmenting interactions as well as in protein stability, but that it is the S1/H1 lid segment that modulates Site 2 binding. In order to access Site 2, Phe199 is repositioned to uncover the mouth of the Site. This movement is predicated on the ability of the adjacent residue, Tyr200, to move in concert with Phe199. In the apo Grp94 NTD, Tyr200 is constrained by the position of H1, so the exposure of Site 2 ultimately depends on relieving the constraints imposed on Tyr200 by H1. The equivalent residues in Hsp90, Phe138 and Tyr139, occupy nearly identical positions to Phe199 and Tyr200 in apo Grp94. Because Site 2 remains closed in Hsp90, this suggests that the constraints imposed on Tyr139 by H1 cannot be relieved. In Grp94, all of the structures of ligand complexes where Site 2 is exposed (Grp94:PU-H54, Grp94:BnIm, Grp94:bme-BnIm, Grp94:PU-H36) ^15,19^ display very large repositioning movements of H1, supporting the notion that the movement of this structural element is important for Site 2 opening.

We can get insight into the origins of the different role S1/H1 plays in the two paralogs by comparing the structures of the unliganded NTDs from Grp94 (PDB code 1yt2) and Hsp90 (PDB code 1yer, 1yes). These structures show that S1 (Grp94 residues 69-78, Hsp90 residues 13-22) is unlikely to play a role in regulating access to Site 2. The structures of S1 from the two paralogs are superimposable, and the five residues that differ between Grp94 and Hsp90 in this region are solvent exposed and make no discriminating interactions. On the other hand, the position of H1 in the two paralogs differs, suggesting that this region is likely to be the discriminating element. The Grp94 and Hsp90 H1s overlap at their N-termini (Grp94 Ala80, Hsp90 Ala24). However, the Grp94 H1 is longer than the Hsp90 H1 by a half turn, and the trajectories of their axes diverge by ∼16 degrees, such that by their C-termini (Grp94 Leu93, Hsp90 Phe37), the two Cα positions are separated by 7.3 Å (**Figure 8a**). This places H1 of Grp94 closer to H4 of the NTD than H1 of Hsp90 is to the equivalent H4 in Hsp90, resulting in a more compact NTD for Hsp90 than for Grp94.

**FIGURE 8.**
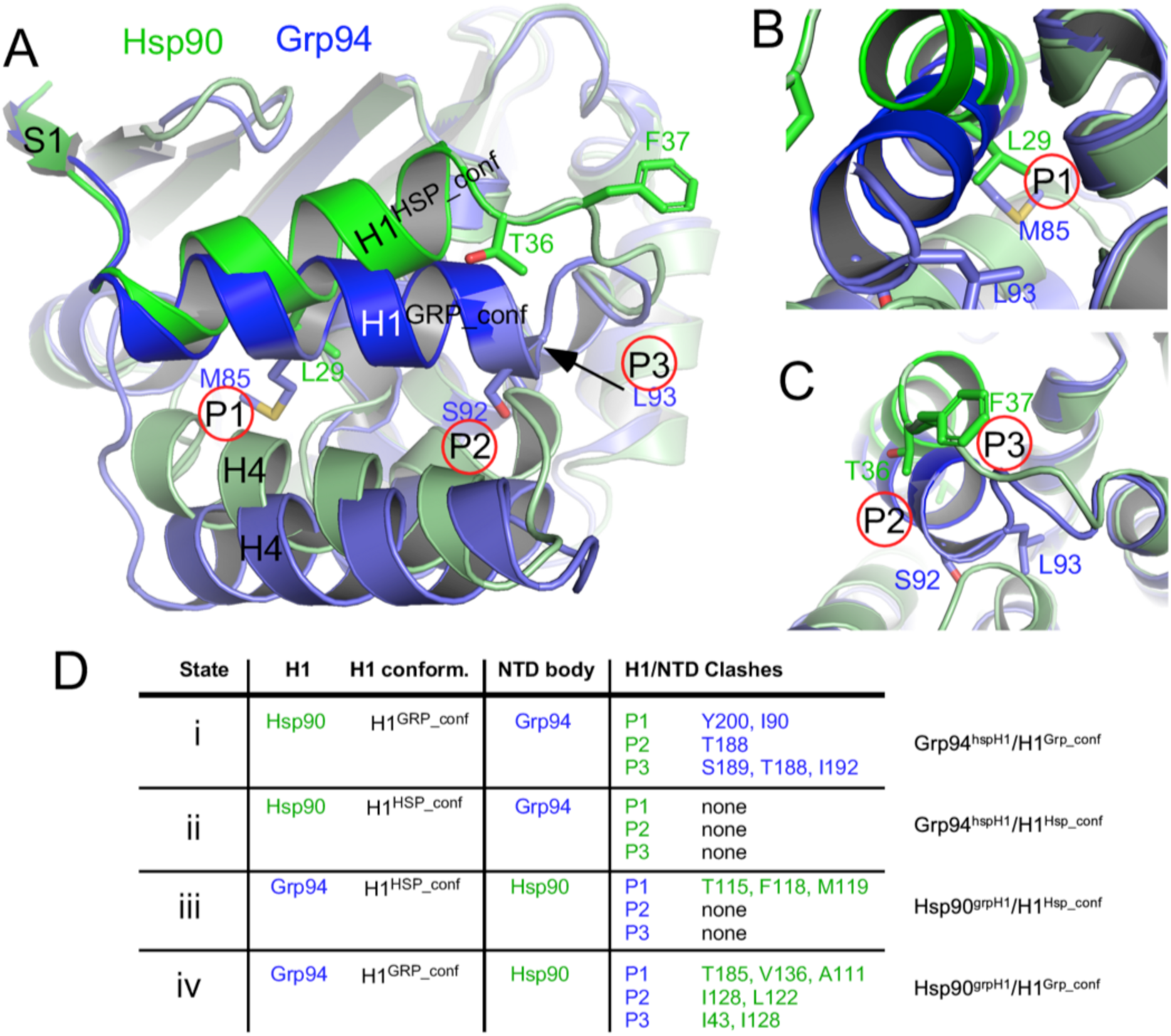
H1 from Hsp90 and Grp94 differ. (A) Overlay of apo Hsp90 and apo Grp94 shows the NTD of Hsp90 is more compact than the NTD of Grp94. The H1 of Hsp90 is closer to the main body of its NTD compared to that of Grp94. The position of H1 in the two paralogs differs such that at their C-termini, the H1s are separated at the Cα by 7.3 Å. The three consequential pairs (P1, P2, P3) of differing residues between the H1s of Hsp90 and Grp94 are shown. Hsp90 is colored green, Grp94 is colored blue. (B) Comparison of the positions of residue pair P1 (Hsp90 L29 and Grp94 M85). (C) Comparison of the positions of residue pairs P2 (Hsp90 T36 and Grp94 S92) and P3 (Hsp90 F37 and Grp94 L93). (D) Summary of modeled interactions in chimeric constructs.

Based on the S1/H1 swap results and by the ∼1.7 °C difference in the T_m_ between Hsp90N and Grp94N presented here (**Figure 6A**), the Hsp90 conformation of H1 (H1^HSP_conf^) is more stable than the Grp94 H1 conformation (H1^GRP_conf^) and less susceptible to movement. We can thus ask what sequence differences permit one paralog to adopt the more stable conformation while the other adopts the less stable position. H1 is comprised of 14 amino acids, of which seven differ between Hsp90 and Grp94 (**Figure 4A**). Four of these differences are likely to be inconsequential due to surface exposure or lack of significant interactions. Three pairs of differing residues, however - Grp94 Met85/Hsp90 Leu29 (P1), Grp94 Ser92/Hsp90 Thr36 (P2), and Grp94 Leu93/Hsp90 Phe37 (P3) – make interactions with, or are closely packed, at the interface with the body of the NTD (**Figure 8A**). To understand whether these three differing pairs of residues can account for the different behaviors of H1, we modeled Grp94 H1 into Hsp90, and Hsp90 H1 into Grp94 using PYMOL. We modeled the swaps in both the stable Hsp90 H1 position (H1^HSP_conf^) and lower stability Grp94 H1 position (H1^GRP_conf^) (**Figure 8D, states i-iv**). From this analysis, numerous steric incompatibilities are apparent.

When H1 of Grp94 is replaced by Hsp90 H1 (Grp94^hspH1^), residues P1, P2, and P3 become Met85Leu, Ser92Thr, and Leu93Phe, respectively. In the first state (Grp94^hspH1^/H1^Grp_conf^) (**Figure 8d, state i**), when the Hsp90 H1 is placed in the more flexible Grp94 position (H1^GRP_conf^), P1, P2, and P3 clash with Grp94 residues Ile90, Tyr200, Thr188, Ser189, and Ile192. These clashes can be completely relieved if the Hsp90 H1 is moved to the stable H1^HSP_conf^ position, showing that Hsp90 H1 is compatible with the body of the Grp94 NTD, but only in the H1^HSP_conf^ position (**Figure 8d, state ii**). Thus, the modeling predicts that Hsp90 H1, when incorporated into Grp94, will adopt the stable H1^HSP_conf^ conformation but not the unstable H1^GRP_conf^ position. This would prevent it from moving in response to Site 2 targeted ligands. This modeling conclusion (Grp94^hspH1^/H1^HSP_conf^) corroborates the experimental data presented above, where the chimeric Grp94^hspS1/H1^ behaved as if Site 2 was no longer accessible to Site 2 binding ligands. The slight increase in T_m_ of Grp94^hspS1/H1^ compared to WT also supports the prediction of the model that H1 is in the H1^HSP_conf^ state (**Figure 6a).**

Conversely, when Grp94 H1 is incorporated into Hsp90 (Hsp90^grpH1^), residues P1, P2, and P3 become Leu29Met, Thr36Ser, and Phe37Leu, respectively. In the opposite of that seen in Grp94^hspH1^, placing the Grp94 H1 of Hsp90^grpH1^ is in the stable position (Hsp90^grpH1^/H1^HSP_conf^), the H1^HSP_conf^ is strongly disfavored because P1 (Leu29Met) would clash with residues Thr115, Phe118, and Met119 of Hsp90 H5 (**Figure 8d, state iii**). The relief of these clashes would require H1 to move away from the body of the NTD, pushing it away from the stable position. This is also in agreement with the experimental data showing that chimeric Hsp90^grpS1/H1^ is accessible to Site 2 binding ligands. Interestingly, if the Grp94 H1 is placed in the more flexible unstable H1^GRP_conf^ position, clashes are also observed between P1, P2, and P3 and Ala111, Thr115, Val136, Ile128, Leu122, Ile43, and Ile128 of the Hsp90 NTD (**Figure 8d, state iv**). This suggests that a different unstable position for Grp94 H1 is achieved in the Hsp90^grpS1/H1^ chimera. The results from the Thermofluor assay described above are consistent with this scenario, where a large decrease of 7.1 °C in the T_m_ of Hsp90^grpS1/H1^ compared to WT Hsp90 is observed. Together, these modeling studies explain how the sequence of Grp94 H1 dictates its placement into a position of lower stability, thus precipitating its movement to expose Site 2 upon selective ligand binding.

The effect of a H1 swap between Grp94 and Hsp90 on the binding of Site 2 selective ligands, and the modeling above suggests that H1 plays a significant role in permitting the remodeling of the ATP binding pocket that drives paralog selective ligand affinity. Site 2 binding is not observed in human Hsp90, but the structure of Hsp90 from *Candida albicans* bound to the fungal-selective ligand CMLD013075 (PDB code 6cjp) ^25^ revealed that CMLD013075 binds to Site 2 and that the position of H1 resembles that seen in Grp94:PU-H36. To accommodate the 4-methoxybenzyl moiety in Site 2, *Candida* Hsp90 Phe127 and Tyr128 (equivalent to Grp94 Phe199 and Tyr200) move away from their pocket-occluding positions seen in the apo structure (PDB code 6cji), thereby precipitating the change in H1 conformation. A comparison of the three H1 residue positions that were identified in the modeling above as determinants of H1 movement shows that while P1 and P2 are identical to their human Hsp90 counterparts, P3 (*Hs*Hsp90 Phe37/*Ca*Hsp90 Val26), differs (**Figure 4b**). In the apo configuration, H1 from human and *Candida* Hsp90 overlap. However, in the CMLD013075-bound structure, H1 is extended at its C-terminus such that Val26 faces H5. Modeling a phenylalanine in place of valine in this conformation predicts a clash with Leu111 of H5. Thus, it appears that P3, the sensitive H1 position in *Candida* Hsp90, allows for movement of H1 by favorably repositioning Val26.

The role of H1 in directing Site 2 access in *Candida* Hsp90 is further supported by the reported mutagenesis data, where a Phe131Tyr mutation in the body of the NTD reduces the sensitivity to CMDL013075. From the structure, the Phe131Tyr mutation appears to stabilize the interaction with Met19 of H1, thereby providing a greater energetic barrier to H1 movement and Site 2 exposure. Finally, it was also noted that CMDL013075 was effective growth suppressor of yeast Hsc82. Yeast Hsc82 also has a valine at P3, the third sensitive H1 position (**Figure 4b)**, supporting the notion that this lowers the barrier to H1 rearrangement in a manner similar to that of *Candida* Hsp90 upon Site 2 binding ligands. Interestingly, a comparison of Hsp90 from multiple organisms ^43^ shows that the incorporation of phenylalanine at P3 is a relatively recent adaptation that is specific to metazoans and higher eukaryotes, suggesting that there is an evolutionary benefit to stiffening the interaction between H1 and the body of the NTD.

In a similar manner, an examination of H1 from Trap1 and *E.coli* HtpG, paralogs that, along with Grp94 are evolutionarily older than eukaryotic Hsp90, indicates that two of the 3 discriminating residues differ from the stable set seen for Hsp90 (**Figure 4b**). In both Trap1 and HtpG, P2 and P3 are Ser-Leu, which is identical to P2 and P3 seen in Grp94. The prediction is that H1 in these proteins would also allow the ATP pocket remodeling to expose Site 2. In support of this, FP competition assays of Grp94-selective PU compounds have consistently showed binding to Trap1 that was significantly stronger than to Hsp90, suggesting that these compounds accessed Site 2 in Trap1 ^15,17^, albeit with a higher energetic penalty than in Grp94.

Recently a novel ATP pocket rearrangement was observed in the structure of human Hsp90 in complex with KUNA-111, an Hsp90-selective, tertiary alcohol inhibitor (PDB code 7ur3) ^44^. In this structure, the side chain of Phe138 adopts a new rotomeric position such that the edge-on-pi interaction with Tyr139 was replaced by a pi-pi interaction with the same residue, thus expanding the volume of Site 1 to accommodate the bulky phenylpropyl substituent. Notably, however, the position of Tyr139 was not perturbed by this new rotomeric position of Phe138, nor was the position or conformation of H1. Modeling of PU-H36 and bme-BnIm into the Hsp90:KUNA-111 structure shows that despite its large rotation, the movement of Phe138 is insufficient to expose Site 2, with steric clashes predicted between the 8-aryl and methylester substituents of these ligands, and the Phe138 beta carbon. Thus, although a novel repositioning of Phe138 has for the first time been observed in human Hsp90, the exposure of Site 2 still appears to require the coordinated movement of the adjacent Tyr139 and H1.

Overall, our results support the key role of H1 in determining the response of the chaperone to ligand binding in the pocket, especially in binding of Site 2 targeted inhibitors. Chaperones with conformationally flexible H1s like Grp94 are sensitive to ligands that target the Site 2 pocket while those with a more rigid H1 attachment to the body of the NTD, like human Hsp90, are impervious and thus preclude the large structural conformational shifts associated with Site 2 binding. H1 can thus be regarded as a selectivity module, and when the relatively immobile H1 of Hsp90 is replaced by the more conformationally flexible equivalent H1 of Grp94, Hsp90 adopts a Grp94-like mode of binding that includes exposing the hydrophobic Site 2 pocket.

Identifying the structural drivers of ligand sensitivity, however, is just one component of solving the puzzle of therapeutic targeting of Grp94. A view is now emerging that multiple structural forms of Grp94 exist in disease, and that these forms are driven in part by post-translational modifications such as *N*-glycosylation and subsequent organization of the chaperone into multi-component epichaperome structures ^45,46^. Further studies will be required to understand how these epigenetic regulatory mechanisms influence the structure of Grp94 and restrict or enhance the access of ligands to selective, targetable side pockets.

Finally, our study has shown that an understanding of what distinguishes the structural characteristics of Hsp90 and its response to ligand from that of Grp94 can greatly benefit from the availability of a chemical tool set, comprised of Grp94-selective and non-selective inhibitors, as well as a library of high resolution structures of Hsp90 and Grp94 in complex with ligands.

## Acknowledgements

Supported by grants R01-CA095130 and P01-CA186866 from the NIH and the Goode Foundation (D.G.). We thank Kevin Maharaj and P.S. for working on an initial but different crystal of Gr94N:PU-H71 and for initial structure refinement. Crystallization screening at the National Crystallization Center at HWI was supported through NIH grant R24GM141256. We thank the NCC personnel, both past and present, for their help. X-ray diffraction data were collected at the Advanced Photon Source beamline 23-IDB, and the Stanford Synchrotron Radiation Lab beamline BL9-2, and initial screening for crystal diffraction was performed at CHESS beamline ID7B2.

## Author contributions

N.Q. designed and performed all experiments with input from D.G. P.S. performed ITC against PU-H54. N.Q. and D.G. analyzed the data. W.A. helped prepare proteins and performed initial ATPase assays. N.Q wrote the original draft. N.Q. and D.G. revised the manuscript.

## References

1. Echeverria PC, Bernthaler A, Dupuis P, Mayer B, Picard D. An interaction network predicted from public data as a discovery tool: application to the Hsp90 molecular chaperone machine. PLoS One. 2011;6(10):e26044.

2. Taipale M, Jarosz DF, Lindquist S. HSP90 at the hub of protein homeostasis: emerging mechanistic insights. Nat Rev Mol Cell Biol. 2010;11(7):515–528.

3. Schopf FH, Biebl MM, Buchner J. The HSP90 chaperone machinery. Nature Reviews Molecular Cell Biology. 2017;18(6):345.

4. Ansa-Addo EA, Thaxton J, Hong F, Wu BX, Zhang Y, Fugle CW, Metelli A, Riesenberg B, Williams K, Gewirth DT, Chiosis G, Liu B, Li Z. Clients and Oncogenic Roles of Molecular Chaperone gp96/grp94. Curr Top Med Chem. 2016;16(25):2765–2778.

5. Gewirth DT. Paralog Specific Hsp90 Inhibitors - A Brief History and a Bright Future. Current topics in medicinal chemistry. 2016;16(25):2779–2791.

6. Valastyan JS, Lindquist S. Mechanisms of protein-folding diseases at a glance. Dis Model Mech. 2014;7(1):9–14.

7. Sanchez J, Carter TR, Cohen MS, Blagg BSJ. Old and New Approaches to Target the Hsp90 Chaperone. Curr Cancer Drug Targets. 2020;20(4):253–270.

8. Dekker FA, Rudiger SGD. The Mitochondrial Hsp90 TRAP1 and Alzheimer’s Disease. Front Mol Biosci. 2021;8:697913.

9. Xie S, Wang X, Gan S, Tang X, Kang X, Zhu S. The Mitochondrial Chaperone TRAP1 as a Candidate Target of Oncotherapy. Front Oncol. 2020;10:585047.

10. Southworth DR, Agard DA. Client-loading conformation of the Hsp90 molecular chaperone revealed in the cryo-EM structure of the human Hsp90:Hop complex. Mol Cell. 2011;42(6):771–781.

11. Dollins DE, Warren JJ, Immormino RM, Gewirth DT. Structures of GRP94-nucleotide complexes reveal mechanistic differences between the hsp90 chaperones. Molecular cell. 2007;28(1):41–56.

12. Huck JD, Que NL, Hong F, Li Z, Gewirth DT. Structural and Functional Analysis of GRP94 in the Closed State Reveals an Essential Role for the Pre-N Domain and a Potential Client-Binding Site. Cell Rep. 2017;20(12):2800–2809.

13. Jhaveri K, Taldone T, Modi S, Chiosis G. Advances in the clinical development of heat shock protein 90 (Hsp90) inhibitors in cancers. Biochimica et biophysica acta. 2012;1823(3):742–755.

14. Yuno A, Lee MJ, Lee S, Tomita Y, Rekhtman D, Moore B, Trepel JB. Clinical Evaluation and Biomarker Profiling of Hsp90 Inhibitors. Methods Mol Biol. 2018;1709:423–441.

15. Patel PD, Yan P, Seidler PM, Patel HJ, Sun W, Yang C, Que NS, Taldone T, Finotti P, Stephani RA, Gewirth DT, Chiosis G. Paralog-selective Hsp90 inhibitors define tumor-specific regulation of HER2. Nat Chem Biol. 2013;9(11):677–684.

16. Taldone T, Patel PD, Patel M, Patel HJ, Evans CE, Rodina A, Ochiana S, Shah SK, Uddin M, Gewirth D, Chiosis G. Experimental and structural testing module to analyze paralogue-specificity and affinity in the Hsp90 inhibitors series. J Med Chem. 2013;56(17):6803–6818.

17. Patel HJ, Patel PD, Ochiana SO, Yan P, Sun W, Patel MR, Shah SK, Tramentozzi E, Brooks J, Bolaender A, Shrestha L, Stephani R, Finotti P, Leifer C, Li Z, Gewirth DT, Taldone T, Chiosis G. Structure-activity relationship in a purine-scaffold compound series with selectivity for the endoplasmic reticulum Hsp90 paralog Grp94. J Med Chem. 2015;58(9):3922–3943.

18. Jiang F, Guo AP, Xu JC, You QD, Xu XL. Discovery of a Potent Grp94 Selective Inhibitor with Anti-Inflammatory Efficacy in a Mouse Model of Ulcerative Colitis. J Med Chem. 2018;61(21):9513–9533.

19. Que NLS, Crowley VM, Duerfeldt AS, Zhao J, Kent CN, Blagg BSJ, Gewirth DT. Structure Based Design of a Grp94-Selective Inhibitor: Exploiting a Key Residue in Grp94 To Optimize Paralog-Selective Binding. J Med Chem. 2018;61(7):2793–2805.

20. Mishra SJ, Khandelwal A, Banerjee M, Balch M, Peng S, Davis RE, Merfeld T, Munthali V, Deng J, Matts RL, Blagg BSJ. Selective Inhibition of the Hsp90alpha Isoform. Angew Chem Int Ed Engl. 2021;60(19):10547–10551.

21. Mishra SJ, Liu W, Beebe K, Banerjee M, Kent CN, Munthali V, Koren J, 3rd, Taylor JA, 3rd, Neckers LM, Holzbeierlein J, Blagg BSJ. The Development of Hsp90beta-Selective Inhibitors to Overcome Detriments Associated with pan-Hsp90 Inhibition. J Med Chem. 2021;64(3):1545–1557.

22. Soldano KL, Jivan A, Nicchitta CV, Gewirth DT. Structure of the N-terminal domain of GRP94. Basis for ligand specificity and regulation. The Journal of biological chemistry. 2003;278(48):48330–48338.

23. Huck JD, Que NLS, Immormino RM, Shrestha L, Taldone T, Chiosis G, Gewirth DT. NECA derivatives exploit the paralog-specific properties of the site 3 side pocket of Grp94, the endoplasmic reticulum Hsp90. The Journal of biological chemistry. 2019;294(44):16010–16019.

24. Immormino RM, Metzger LEt, Reardon PN, Dollins DE, Blagg BS, Gewirth DT. Different poses for ligand and chaperone in inhibitor-bound Hsp90 and GRP94: implications for paralog-specific drug design. Journal of molecular biology. 2009;388(5):1033–1042.

25. Whitesell L, Robbins N, Huang DS, McLellan CA, Shekhar-Guturja T, LeBlanc EV, Nation CS, Hui R, Hutchinson A, Collins C, Chatterjee S, Trilles R, Xie JL, Krysan DJ, Lindquist S, Porco JA, Jr., Tatu U, Brown LE, Pizarro J, Cowen LE. Structural basis for species-selective targeting of Hsp90 in a pathogenic fungus. Nat Commun. 2019;10(1):402.

26. He H, Zatorska D, Kim J, Aguirre J, Llauger L, She Y, Wu N, Immormino RM, Gewirth DT, Chiosis G. Identification of potent water soluble purine-scaffold inhibitors of the heat shock protein 90. J Med Chem. 2006;49(1):381–390.

27. Luft JR, Collins RJ, Fehrman NA, Lauricella AM, Veatch CK, DeTitta GT. A deliberate approach to screening for initial crystallization conditions of biological macromolecules. J Struct Biol. 2003;142(1):170–179.

28. Otwinowski Z, Minor W. Processing of X-ray diffraction data collected in oscillation mode. Methods in Enzymology. 1997;276:307–326.

29. Vonrhein C, Flensburg C, Keller P, Sharff A, Smart O, Paciorek W, Womack T, Bricogne G. Data processing and analysis with the autoPROC toolbox. Acta crystallographica. 2011;67(Pt 4):293–302.

30. STARANISO [computer program]. Cambridge, United Kingdom: Global Phasing Ltd.; 2018.

31. Liebschner D, Afonine PV, Baker ML, Bunkoczi G, Chen VB, Croll TI, Hintze B, Hung LW, Jain S, McCoy AJ, Moriarty NW, Oeffner RD, Poon BK, Prisant MG, Read RJ, Richardson JS, Richardson DC, Sammito MD, Sobolev OV, Stockwell DH, Terwilliger TC, Urzhumtsev AG, Videau LL, Williams CJ, Adams PD. Macromolecular structure determination using X-rays, neutrons and electrons: recent developments in Phenix. Acta Crystallogr D Struct Biol. 2019;75(Pt 10):861–877.

32. Emsley P, Cowtan K. Coot: model-building tools for molecular graphics. Acta crystallographica. 2004;60(Pt 12 Pt 1):2126–2132.

33. Schuttelkopf AW, van Aalten DM. PRODRG: a tool for high-throughput crystallography of protein-ligand complexes. Acta Crystallogr Sect D Biol Crystallogr. 2004;60(Pt 8):1355–1363.

34. Chen VB, Arendall WB, 3rd, Headd JJ, Keedy DA, Immormino RM, Kapral GJ, Murray LW, Richardson JS, Richardson DC. MolProbity: all-atom structure validation for macromolecular crystallography. Acta Crystallogr Sect D Biol Crystallogr. 2010;66(Pt 1):12–21.

35. Immormino RM, Kang Y, Chiosis G, Gewirth DT. Structural and quantum chemical studies of 8-aryl-sulfanyl adenine class Hsp90 inhibitors. J Med Chem. 2006;49(16):4953–4960.

36. Ernst JT, Liu M, Zuccola H, Neubert T, Beaumont K, Turnbull A, Kallel A, Vought B, Stamos D. Correlation between chemotype-dependent binding conformations of HSP90alpha/beta and isoform selectivity-implications for the structure-based design of HSP90alpha/beta selective inhibitors for treating neurodegenerative diseases. Bioorg Med Chem Lett. 2014;24(1):204–208.

37. Austin C, Pettit SN, Magnolo SK, Sanvoisin J, Chen W, Wood SP, Freeman LD, Pengelly RJ, Hughes DE. Fragment screening using capillary electrophoresis (CEfrag) for hit identification of heat shock protein 90 ATPase inhibitors. J Biomol Screen. 2012;17(7):868–876.

38. Wright L, Barril X, Dymock B, Sheridan L, Surgenor A, Beswick M, Drysdale M, Collier A, Massey A, Davies N, Fink A, Fromont C, Aherne W, Boxall K, Sharp S, Workman P, Hubbard RE. Structure-activity relationships in purine-based inhibitor binding to HSP90 isoforms. Chem Biol. 2004;11(6):775–785.

39. Sigurskjold BW. Exact analysis of competition ligand binding by displacement isothermal titration calorimetry. Anal Biochem. 2000;277(2):260–266.

40. Velazquez-Campoy A, Freire E. Isothermal titration calorimetry to determine association constants for high-affinity ligands. Nat Protoc. 2006;1(1):186–191.

41. Krukenberg KA, Street TO, Lavery LA, Agard DA. Conformational dynamics of the molecular chaperone Hsp90. Quarterly reviews of biophysics. 2011;44(2):229–255.

42. Roher N, Miro F, Boldyreff B, Llorens F, Plana M, Issinger OG, Itarte E. The C-terminal domain of human grp94 protects the catalytic subunit of protein kinase CK2 (CK2alpha) against thermal aggregation. Role of disulfide bonds. Eur J Biochem. 2001;268(2):429–436.

43. Starr TN, Flynn JM, Mishra P, Bolon DNA, Thornton JW. Pervasive contingency and entrenchment in a billion years of Hsp90 evolution. Proceedings of the National Academy of Sciences of the United States of America. 2018;115(17):4453–4458.

44. Mishra SJ, Reynolds TS, Merfeld T, Balch M, Peng S, Deng J, Matts R, Blagg BSJ. Structure-Activity Relationship Study of Tertiary Alcohol Hsp90alpha-Selective Inhibitors with Novel Binding Mode. ACS Med Chem Lett. 2022;13(12):1870–1878.

45. Rodina A, Wang T, Yan P, Gomes ED, Dunphy MP, Pillarsetty N, Koren J, Gerecitano JF, Taldone T, Zong H, Caldas-Lopes E, Alpaugh M, Corben A, Riolo M, Beattie B, Pressl C, Peter RI, Xu C, Trondl R, Patel HJ, Shimizu F, Bolaender A, Yang C, Panchal P, Farooq MF, Kishinevsky S, Modi S, Lin O, Chu F, Patil S, Erdjument-Bromage H, Zanzonico P, Hudis C, Studer L, Roboz GJ, Cesarman E, Cerchietti L, Levine R, Melnick A, Larson SM, Lewis JS, Guzman ML, Chiosis G. The epichaperome is an integrated chaperome network that facilitates tumour survival. Nature. 2016;538(7625):397-401.

46. Yan P, Patel HJ, Sharma S, Corben A, Wang T, Panchal P, Yang C, Sun W, Araujo TL, Rodina A, Joshi S, Robzyk K, Gandu S, White JR, de Stanchina E, Modi S, Janjigian YY, Hill EG, Liu B, Erdjument-Bromage H, Neubert TA, Que NLS, Li Z, Gewirth DT, Taldone T, Chiosis G. Molecular Stressors Engender Protein Connectivity Dysfunction through Aberrant N-Glycosylation of a Chaperone. Cell Rep. 2020;31(13):107840.

