## Supplemental Figures for "Selective inhibition of hsp90 paralogs: Uncovering the role of helix 1 in Grp94-selective ligand binding"

### Supplementary Data

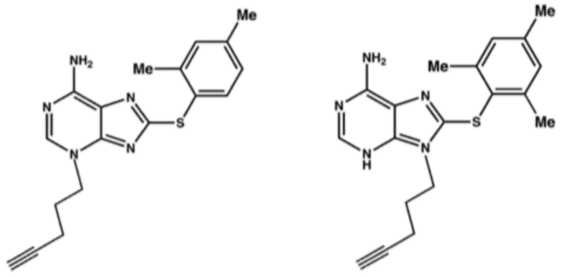

PU-H54 PU-H36

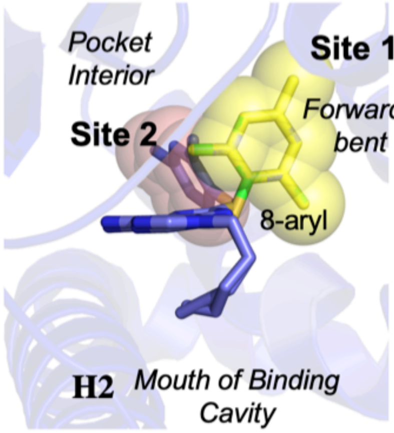


**FIGURE S1**. **The pose of PU-H54 structure in the binding pocket depends on which accessory site binds the 8-aryl moiety**. In Hsp90, the 8-aryl moiety is found in a forward bent conformation as it is inserted into Site 1. In Grp94 the 8-aryl group is in Site 2 and thus adopts a backward bent pose. The inhibitor PU-H36 differs from PU-H54 in having its acetenlynic tail at N9 instead of N3 as well as having a more trimethylated 8-aryl group.

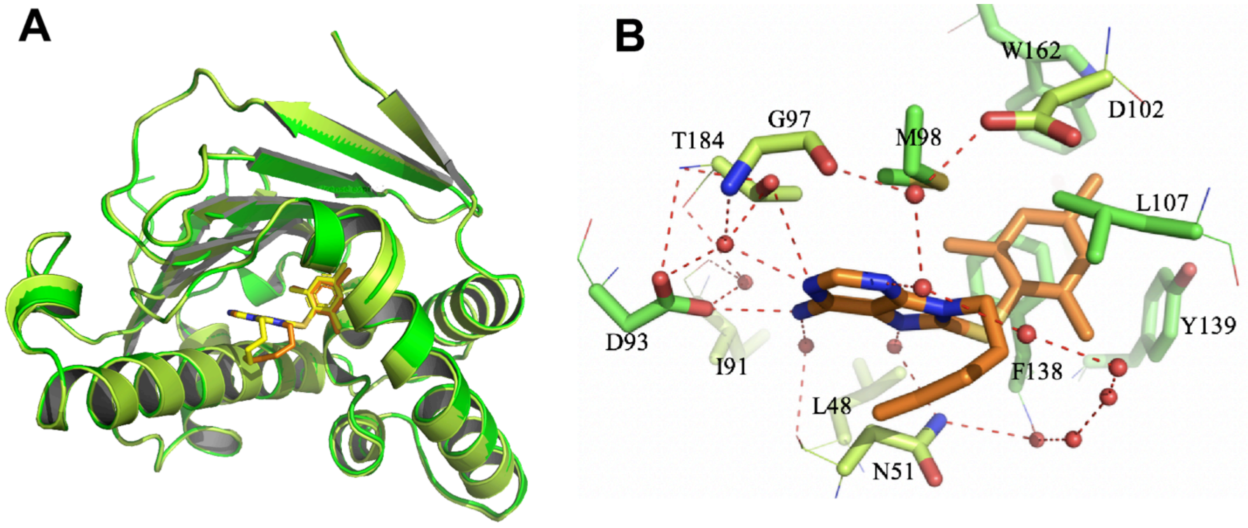


**FIGURE S2. PU-H36 binds to Site 1 in Hsp90**.
(A) Superposition of Hsp90:PU-H36 (dark green, orange ligand) with Hsp90:PU-H54 (light green, yellow ligand). In both cases the 8-aryl moiety is in Site 1 and the alkyl tails both adopt the same “scorpion” tail conformation.
(B) PU-H36 interactions in the ATP binding pocket. The 8-aryl moiety in Site 1 makes pi-pi interactions with Phe138 and Tyr139 and van der Waals contact with Leu107. The 5’-methyl sits above the adenine ring and forms a van der Waals contact. Leu107 makes a hydrophobic interaction with the 8-aryl ring.

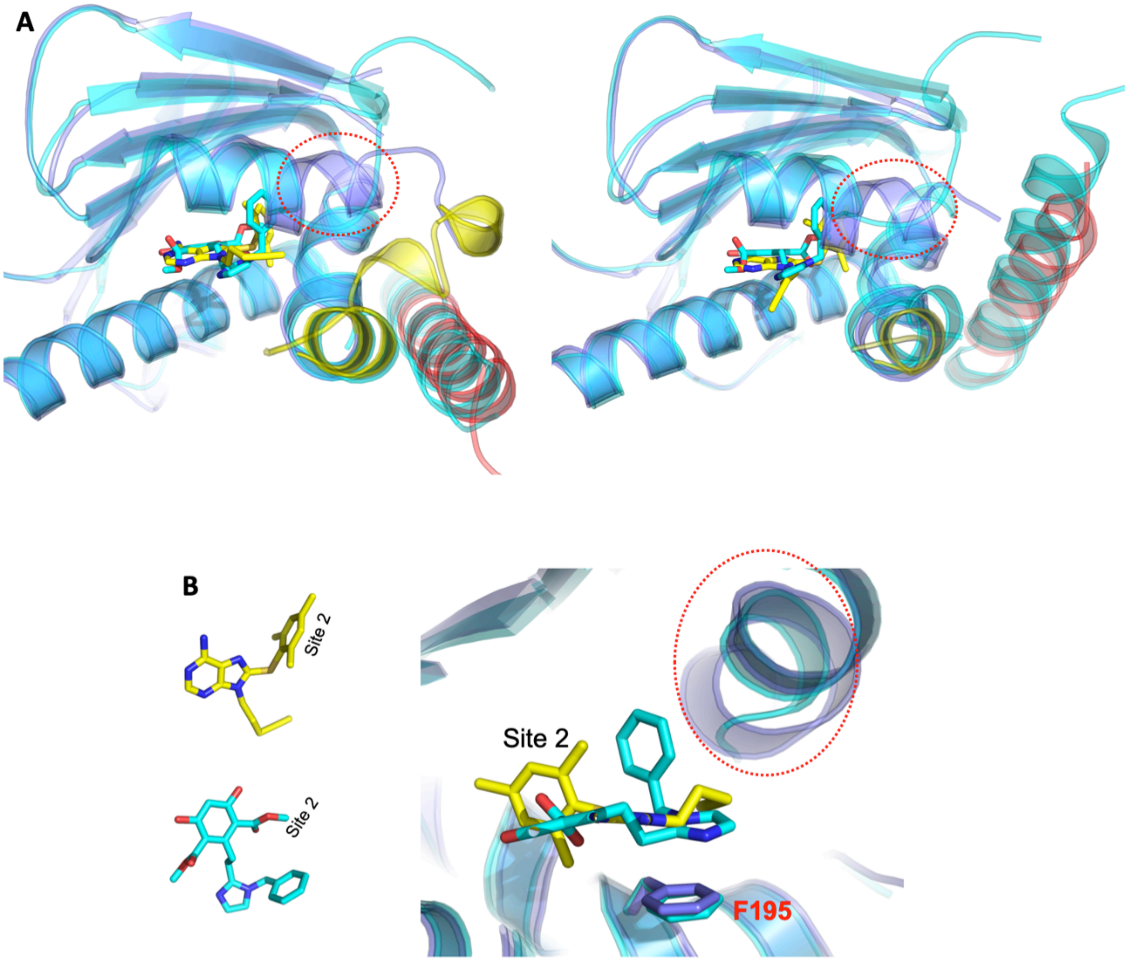


**FIGURE S3**. **The structure of Grp94N:PU-H36 is similar to that of Grp94N: bme-BnIm**. (A) The two ligands can access Site 2 of Grp94 and induce the same conformation of helix 1 in the two monomers comprising the crystallographic subunit. Grp94N:PU-H36 (slate), helix 1 in red, helix 4,5 in yellow; bme-BnIm (cyan) (PDB 6BAW).

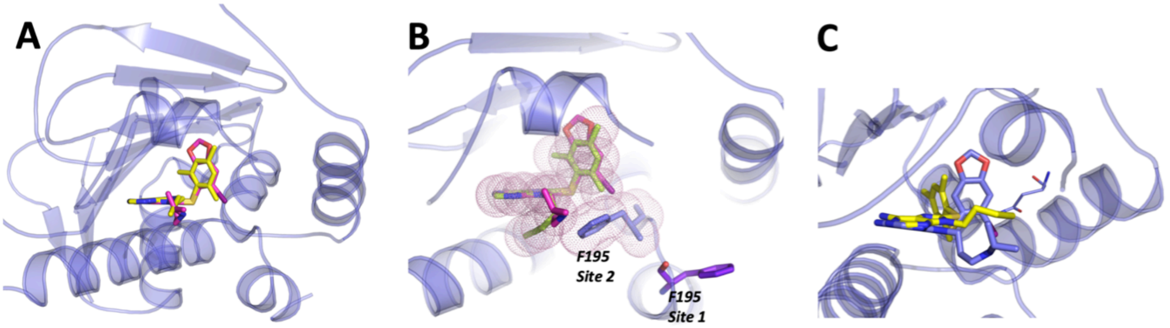


**FIGURE S4**. **PU-H36 modeled in Site 1 of Grp94$^{HspS1/H1}$**.

(A) PU-H36 (yellow) in its forward bent pose when it is bound into Site 1 is modeled into the Grp94N:PU-H71 structure. The 8-aryl of PU-H36 can be inserted into Site 1 similar to that of PU-H71 (magenta).

(B) The Site 1-bound pose of PU-H36 clashes with the side chain of Phe195 in the Site 2 occupied Grp94:PU-H36 structure. Thus F195 and Helix 5 of Grp94$^{HspS1/H1}$ most likely assume the conformation of Helix 5 found in Grp94:PU-H71.

(C) in this model, the alky tail of PU-H36 most likely assumes a pose similar to that of PU-H36 in Grp94:PU-H36 molecule C and PU-H71 in Grp94:PU-H71 or the scorpion tail found in Hsp90:PU-H36. A steric clash between the alkyl tail in the pose shown in Grp94N:PU-H36 molecule A and N162 when it is in the Grp94:PU-H71 conformation. PU-H71 in blue. All models here are by PYMOL.

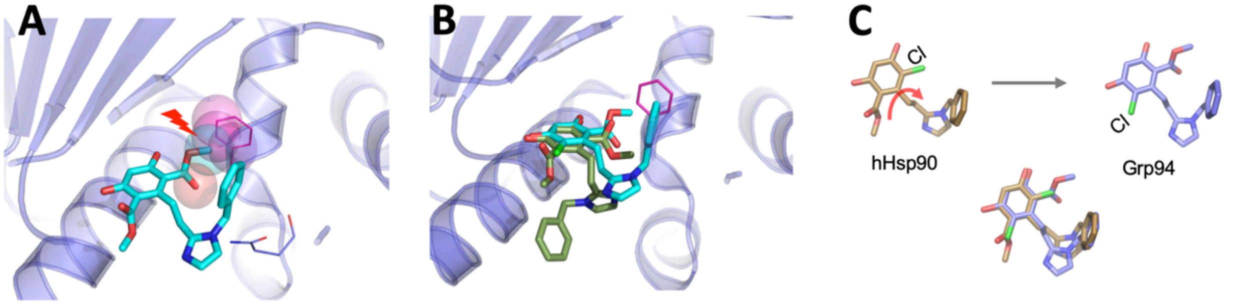


**FIGURE S5**. **The two methyl esters of the resorcinol ring in bme-BnIm makes it impossible to adopt an alternate pose unlike that of inhibitor BnIm**.
(A) The methyl ester of bme-BnIm and its steric clash with the side chain of Phe199 makes it difficult to be stabilized in Site 1 of the Grp94 as shown when bme-BnIm is modeled into the Grp94:PU-H71 structure. The imidazole also shows a steric clash with the Asn162 side chain in this conformation.
(B) The resorcinol of bme-BnIm from PDB 6ceo is not as deeply inserted into the ATP cavity and its pendant moiety is in altered conformation.
(C) The resorcinol ring of BnIm has a chlorine atom instead of a methylester at the 3-position. By flipping this ring about the resorcinol-imidazole linker, the ligand can bind to either Hsp90 or to Grp94. In Grp94 the methyl ester projects into Site 2 and the chlorine is oriented towards helix 2. Because Hsp90 cannot open up Site 2, the ring is flipped such that the chlorine atom is projected towards helix 6 and does not mediate Site 2 binding. All models here are by PYMOL

```
hTrap1       173  GTIARSGSKAFLDALQNQA---EASSKIIGQFGVGFYSAFMVAD  213
HTPG          95  GTIAKSGTKSFLESLGSDQ---AKDSQLIGQFGVGFYSAFIVAD  135
Grp94        164  GTIAKSGTSEFLNKMTEAQEDGQSTSELIGQFGVGFYSAFLVAD  207
hHsp90A      108  GTIAKSGTKAFMEALQAG-----ADISMIGQFGVGFYSAYLVAE  146
hHsp90B      103  GTIAKSGTKAFMEALQAG-----ADISMIGQFGVGFYSAYLVAE  141
FungHsp90     97  GTIAKSGTKSFMEALSAG-----ADVSMIGQFGVGFYSLFLVAD  135
yhsp82        94  GTIAKSGTKAFMEALSAG-----ADVSMIGQFGVGFYSLFLVAD  132
yhsc82        94  GTIAKSGTKAFMEALSAG-----ADVSMIGQFGVGFYSLFLVAD  132
```

**FIGURE S6. Helix 4/Helix 5 of Grp94 and Hsp90**. The sequence in blue was swapped between Hsp90 and Grp94. Sequence alignment of the Helix 4/Helix 5 regions of various hsp90s.
